# The mRNA architecture of the termination site primes programmed stop codon readthrough events in *Drosophila*

**DOI:** 10.1101/2025.11.07.685291

**Authors:** Laveena Kansara, Georg Wolfstetter, Daniel Wintermayr, Ioannis Alexopoulos, Alejandra Escós, Marc R. Friedländer, Ylva Engström

## Abstract

Programmed stop codon readthrough (SCR) is a form of genetic re-coding, in which a near-cognate tRNA base-pairs with a stop codon, leading to the translation of a C-terminally extended protein. Recent studies revealed that SCR represents an evolutionarily conserved, spatio-temporally controlled mechanism of posttranscriptional gene regulation that requires *cis*-regulatory elements as well as *trans*-acting factors. In this study, we characterized cis-regulatory elements controlling programmed SCR of the *Drosophila* POU3-family member *drifter/ventral veins lacking* (*dfr/vvl*). Using S2 cell-based luciferase assays, we show that stop codon identity and the +4 to +9 nucleotide sequence are required but not sufficient for *dfr* SCR regulation. Phylogenetic prediction identified an mRNA stem-loop in the 3’ UTR, proximal to the readthrough UAG codon. Mutational analysis revealed that the distance from the stop codon as well as stem-loop stability, but not the underlying sequence identity, critically impact *dfr* SCR. Similarly, the mRNA stem-loop promoted SCR in an *in vivo Drosophila* model. We applied this information to refine computational prediction of SCR-associated mRNA stem-loops and show that these elements effectively promote SCR of heterologous mRNAs. These findings increase our understanding of SCR and the underlying regulatory mechanisms.

**Key Points:**

- Stop codon readthrough is regulated both by mRNA sequence identity and mRNA structure elements.
- A 3’ mRNA stem-loop and its thermodynamic stability determine stop codon readthrough efficiency.
- mRNA stem-loops are frequently found in Drosophila genes that exhibit stop codon readthrough.

**Graphical Abstract:** 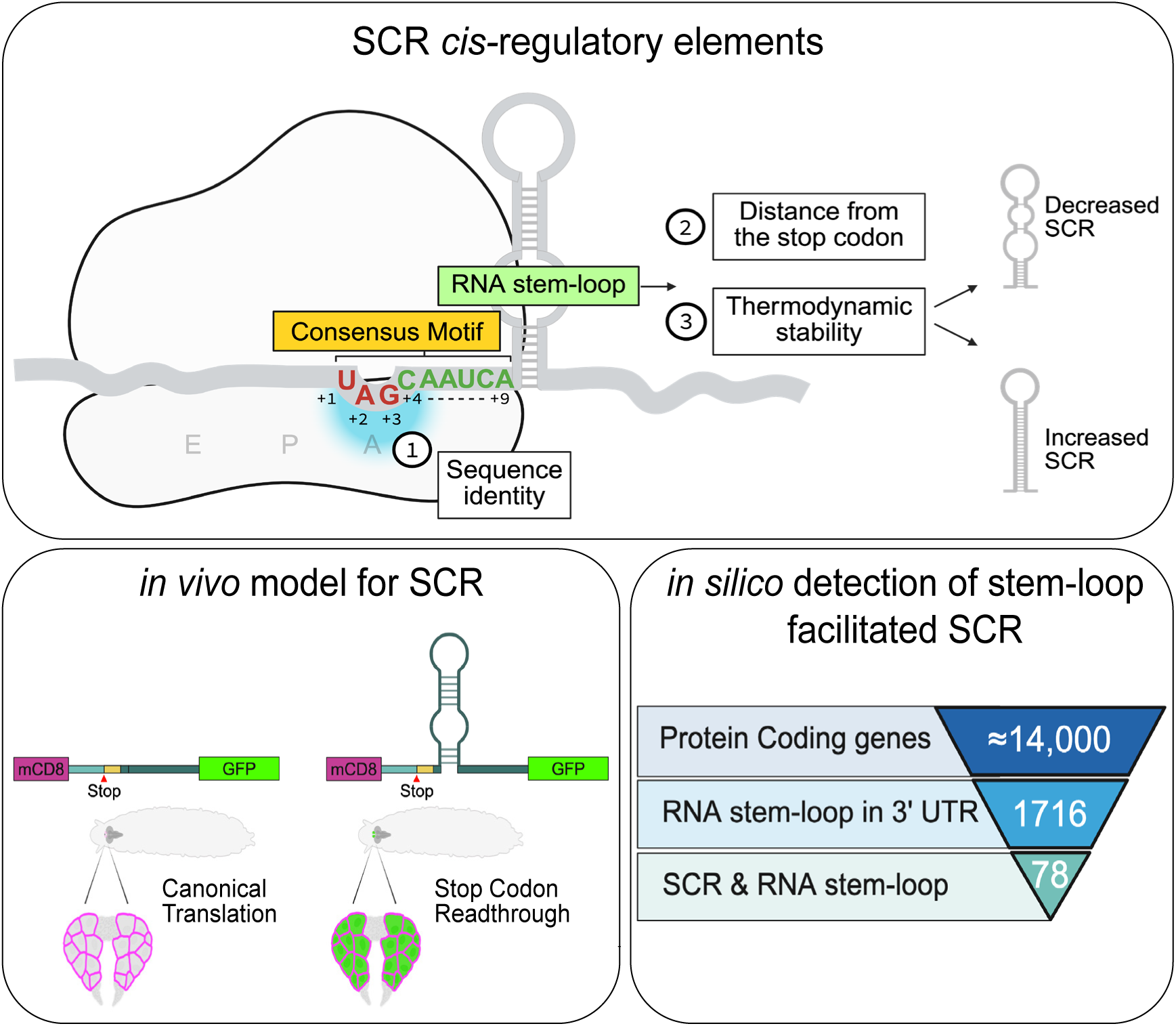

## Introduction

Canonical translation of mRNAs usually terminates when the ribosome encounters a stop codon, however, translational termination can be bypassed in non-standard decoding events like ribosome hopping, frame-shifting, and by re-programming of the stop codon message ^1, 2^. The stop codon message can be re-programmed by the incorporation of a near-cognate tRNA to the ribosome’s A-site, which competes with the recruitment of the translation termination complex. This mechanism, referred to as programmed stop codon readthrough (SCR), was initially studied in certain retro- and RNA-viruses that exploit SCR in their hosts to increase genome coding capacities ^3^. Other exceptions have been described in bacteria and yeast where stop codons can be re-programed by nonsense suppressor tRNAs in order to increase proteome diversity or to adapt to environmental conditions ^4^. Another extensively studied re-coding mechanism is the incorporation of selenocysteine at the position of an UGA stop codon, which is promoted by selenocysteine insertion sequence (SECIS) mRNA stem loops ^5^.

Recent *in silico* predictions and ribosome profiling approaches have uncovered wide-spread SCR-events in fungi, animals, and plants. It has now become evident that programmed SCR represents a highly evolutionarily conserved, spatio-temporally controlled process that increases proteome diversity ^6-8^. Comparative phylogenetic analyses in insects have revealed the existence of conserved open reading frames (ORFs) in the 3’ UTR of several hundred genes ^9-11^. However, the overlap between these phylogenetic predictions and data from independent ribosome profiling studies has been unsatisfactory ^12^. Moreover, there remains a large gap between the number of predicted SCR events and the few instances in which SCR has been functionally characterized. Thus, our knowledge about the underlying mechanisms and functional consequences remain scarce.

In *Drosophila melanogaster*, spatio-temporally controlled SCR has been experimentally demonstrated for the genes *headcase (hdc),* the founding homolog of the vertebrate *HECA* genes ^13^, *kelch (kel)* encoding a substrate-targeting protein associated with the Cullin3-RING ubiquitin E3 ligase complex ^14, 15^, *traffic jam (tj)* encoding a MAF-like bZIP transcription factor ^16^; *Synapsin* ^17^; and the *POU3-family* transcription factor homolog *drifter/ventral veins lacking (dfr/vvl),* hereafter referred to as *dfr* ^18^. *dfr* is an evolutionarily conserved, intron-less gene that encodes a ∼45.9 kDa protein (referred to as the canonical short Dfr isoform, or Dfr-S). Notably, the first 858 bp of the *dfr* 3’ untranslated region (3’ UTR) exhibit a high degree of sequence conservation amongst dipteran species and comprise a consecutive codon message in frame to Dfr-S. Consequently, *dfr* mRNA has been predicted as a likely candidate to undergo programmed SCR ^11^. This was experimentally confirmed by Zhao et al. ^18^. revealing tissue-specific, post-embryonic expression of a C-terminally extended ∼77 kDa long Dfr isoform (Dfr-L), in which the UAG stop codon is re-coded to glutamine. Further functional analyses uncovered the involvement of Dfr-L in organismal growth, steroid hormone biogenesis and immune defense while the canonical Dfr-S has additional essential roles during embryonic development ^18-20.^

Previous studies suggest that a combination of *cis-* and *trans-*acting elements might be required to facilitate programmed stop codon readthrough ^6^. For instance, the nucleotide sequence surrounding the read-through stop codon exhibits a remarkable level of evolutionary conservation ^21, 22^ most likely reflecting a requirement for direct interactions with the translation machinery. Moreover, studies have predicted the presence of mRNA secondary structures in the 3’ proximity of the stop codon that are distinct from the well-characterized SECIS stem loops and represent a predictive feature of genes undergoing programmed SCR. Previous analyses of the *Drosophila hdc* and *kel* loci experimentally linked 3’ UTR regions to programmed SCR, but although these were predicted to form mRNA structures it remains unclear if it involves secondary structures or other features provided by the mRNA sequence context ^13, 15^. So far, a direct connection between mRNA structure and SCR has only been experimentally confirmed for viral transcripts ^23, 24^. In the present study, we investigated molecular mechanisms regulating programmed SCR, using the *Drosophila dfr* gene as a target. Several *cis*-acting elements that control the rate of SCR were identified in the 3’ UTR of *Dfr* mRNA. The downstream, proximal mRNA stem-loop structure acted as an SCR-promoting element of all the three stop codons with different efficiency. Notably, the structural stability of the stem-loop and the precise distance from the stop codon, but not the mRNA sequence itself, correlated with SCR efficiency. The importance of the mRNA stem-loop for SCR was further confirmed in transgenic *Drosophila* larvae. This information was then used to design genome-wide structure predictions that identified 68, previously unrecognized mRNA secondary structures, located in the 3’ UTR of predicted SCR genes. The experimental validation of six of these supports our model, wherein mRNA stem-loops situated in the proximal 3’ context of a SCR-prone stop codon function as a *cis*-element to facilitate programmed SCR.

## Results

### Accessing translational readthrough of *dfr* mRNA

To identify *cis*-acting elements that regulate *dfr* SCR, we established a two-plasmid luciferase reporter assay in *Drosophila* S2 cells, with Renilla luciferase (RLuc) as a ratio-metric control and Firefly luciferase (FLuc) activity to measure SCR (Fig. 1a). A reporter construct, spanning the translated regions of the *dfr* gene; translon 1 (ORF 1) and translon 2 (ORF 1+2) with translation extending over the first annotated UAG stop codon (*dfr Full*, Fig. 1b), conferred moderate SCR levels, measured as relative FLuc/RLuc luminescence (Fig. 1c). Noteworthy, it has previously been shown that there is neither mRNA editing nor splicing of the stop codon of *dfr,* which is an intron-less gene ^18^. Deletion of the *dfr* C-terminal extension sequence (*dfrΔORF2*) completely abolished FLuc activity, indicating that sequences downstream of the stop codon are essential for *dfr* translational readthrough (Fig. 1b,c). To further map the SCR-promoting region, we analyzed a series of deletion constructs (Fig. 1b). The highest readthrough rates were observed with *dfr288*, which contains the region from -99 nt to +189 relative to the stop codon. Shortening of ORF2 to +39 in *dfr138* significantly decreased the SCR levels compared to *dfr288*, demarcating a positive regulatory element between +39 to +189 (Fig. 1d). However, *dfr138* still promoted a significant level of SCR, to a similar extent as the *dfrFull* construct (Fig. 1c), suggesting that the most proximal region promotes basal levels of SCR (Fig. 1d). RT-qPCR confirmed that the differences in SCR levels across various test cassettes were not caused by major changes in transcript stability (Supp. Fig. S1).

**Figure 1:**
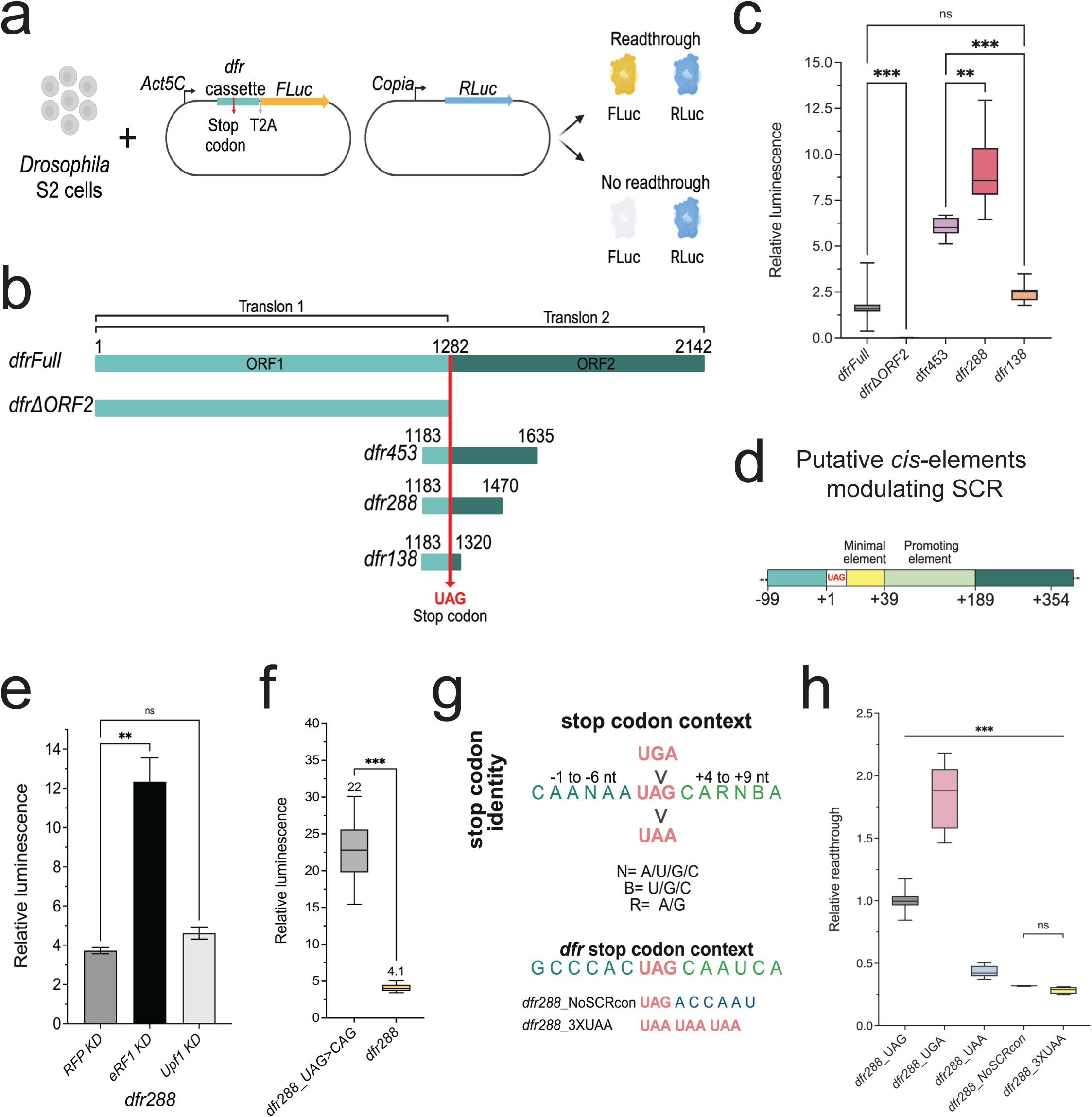
Identification of sequence requirements for *dfr* mRNA readthrough. **a** Illustration of the luciferase reporter assay created for characterization of sequence elements involved in the regulation of SCR. **b** Schematic representation of the test cassettes from the *dfr* gene inserted upstream of FLuc in the reporter plasmid. **c** Box plot diagram depicting relative luminescence (FLuc/RLuc) of *Drosophila* S2 cell samples transfected with the *dfr* constructs indicated on the X-axis. Statistical significance for all readthrough reporter assays was tested by one-way ANOVA with Dunnett’s T3 multiple comparisons test, unless otherwise stated (*n =* 6 independent experimental replicates, *** = *p* ≤ *0.001,* **=*p*= 0.002, *ns= 0.05*). In all box plot diagrams, boxes represent the 25-75% quartile, lines represent the median, whiskers represent min to max. **d** Illustration depicting the *cis*-acting elements identified to be involved in the regulation of *dfr* SCR. Schematics were created with BioRender.com **e** Column graph depicting the relative luminescence values of cells transfected with the *dfr288* construct, dsRNA treatment is represented on the X-axis as knock-down (KD) of mRNAs for *Upf1*, *eRF1*, and control *RFP*. (Data represent the mean ± SEM of three independent experimental replicates (*n=3),* **=*p*= 0.002, *ns= 0.05*). **f** Box plot diagram of relative luminescence in *Drosophila* S2 cell extracts transfected with *dfr288* and the sense codon variant *dfr288_UAG>CAG*. Mean values are indicated above the whiskers. (*n =* 3 independent experimental replicates, *** = *p* ≤ *0.001*). **g** Schematic comparison of the consensus sequence for high SCR. The orientation of stop codons from top to bottom represents their readthrough efficiency, with UGA being the most efficiently suppressed stop and UAA being the least. The sequence surrounding the stop represents the SCR consensus sequence identified from naturally occurring SCR events ^6^. The corresponding region in the *dfr* gene is represented at the bottom. **h** Box plot diagram depicting Relative Readthrough Efficiency (see methods for description) observed after transfection with the indicated *dfr288*_variants: *dfr288_UGA* (UAG to UGA), *dfr288_UAA* (UAG to UAA), *dfr288_NoSCRcon* (+4 to +9nt sequence changed to a non-optimal sequence-CAA UCA>ACC AAU), and *dfr288_+9A>C* (single nucleotide substitution at +9 position). (*n =* 3 independent experimental replicates, *** = *p* ≤ *0.001, ns= 0.05*).

Based on the high SCR levels observed with the *dfr288* reporter, we decided to use this construct for the further characterization of *cis-*acting elements. To confirm that *dfr288* constitutes a valid reporter for suppression of termination, it was tested in combination with RNAi-mediated knock-down of the *eukaryotic release factor 1* (*eRF1*), which led to a 3-fold increase in *dfr288* SCR levels (Fig. 1e). On the contrary, knock-down of *up-frameshift suppressor 1 homolog* (*Upf1*), a key component in the nonsense-mediated mRNA decay (NMD) pathway, did not significantly alter *dfr288* SCR levels (Fig. 1e). Thus, *dfr288* transcripts evade NMD and *dfr288 SCR* provide a quantitative read-out of termination suppression. Finally, following the MINDR guidelines for SCR measurements ^25^ we created a sense codon variant of *dfr288* (*dfr288_UAG>CAG*), which showed about 18% readthrough efficiency for *dfr288* in S2 cells (Fig. 1f). This sense codon control then served as the reference for calculating readthrough efficiency in the following experiments.

Taken together, we conclude that sequences necessary for basal *dfr* SCR are located within a -99 to +39 nt region relative to the stop codon (SCR minimal element, Fig. 1d). A positive *cis-*acting element was mapped between nt +39 to +189 (SCR promoting element, Fig. 1d), which significantly stimulated SCR beyond basal levels, while sequence extensions further upstream of -99 and downstream of +189 reduced FLuc activity.

### The identity of the stop codon and sequences downstream significantly impact *dfr* translational readthrough

The *dfr* mRNA contains a UAG stop codon and a +4 to +9 nucleotide sequence that closely resembles the reported consensus motifs (Fig. 1g) for translational readthrough ^26, 27^. However, the nucleotide sequence from positions -6 to -1 upstream of the stop codon differs significantly from previously reported SCR consensus ^28, 29^. To investigate if the stop codon identity influences readthrough of *dfr*, we changed the UAG codon to UGA and UAA in *dfr288* (Fig. 1h). In agreement with previous findings ^16, 30^, the UAA codon was the most effective termination codon (low SCR) (Fig. 1h), followed by UAG, while UGA was the most efficiently suppressed stop codon, promoting highest readthrough rates. The transcript levels for the stop codon variants remained comparable, indicating that the differences in SCR levels are due to translational regulation and not to mRNA stability (Supp. Fig. S1). In addition to the stop codon identity, the +4 nt to +9 nt sequence context was crucial for *dfr*-FLuc translational readthrough. After changing the block of +4 nt to +9 nt to a non-SCR-favorable ACCAAU sequence (*dfr288_NoSCRcon,* Fig. 1g), we observed significantly reduced FLuc activity (Fig. 1h, Supp. Fig. S1). In addition, swapping the *dfr* stop codon and the +4 to +9 nt block with a triple UAA stop codon sequence (*dfr288*_3xUAA, Fig. 1g) led to a similar drop in readthrough levels (Fig. 1h), suggesting a combined effect of stop codon identity and downstream sequence in SCR regulation.

Thus, our findings corroborate previous observations that stop codon identity and the +4 nt to +9 nt sequence context represent some of the key features of genes prone to exhibit SCR ^26-31^. We further conclude that the stop codon identity and proximal sequences in *dfr* constitute an essential minimal SCR-enabling element (Fig. 1d). However, these sequences alone are insufficient to drive high-level readthrough, which in addition requires sequences further 3’, constituting an SCR promoting element (Fig. 1d).

### An mRNA secondary structure element mediates *dfr* mRNA translational readthrough

To investigate the evolutionary constraints acting on the 3’ UTR and to identify highly conserved sequences that may confer regulatory functions we conducted a Multiple Sequence Alignment (MSA) of the *dfr* gene using 29 *Drosophila* species and two outgroup dipterans (Fig. 2a, Supp. Fig. S2, Supp. Table ST1). The analysis revealed a remarkably high conservation score at the nucleotide level from -18 to +68 region, indicating strong selective pressure to maintain the exact nucleotide sequence (Fig. 2a). In contrast, sequences downstream of this conserved segment displayed the typical pattern of synonymous substitutions at the 3^rd^ nt position (Fig. 2a, Supp. Data SD1), indicating evolutionary conservation at the amino acid coding level, but not at the nucleotide level ^18^. This extensive sequence conservation around the stop codon and to +68 nt suggests the existence of functional *cis*-acting element(s), either at the mRNA sequence or mRNA structure level, or a combination of both.

**Figure 2:**
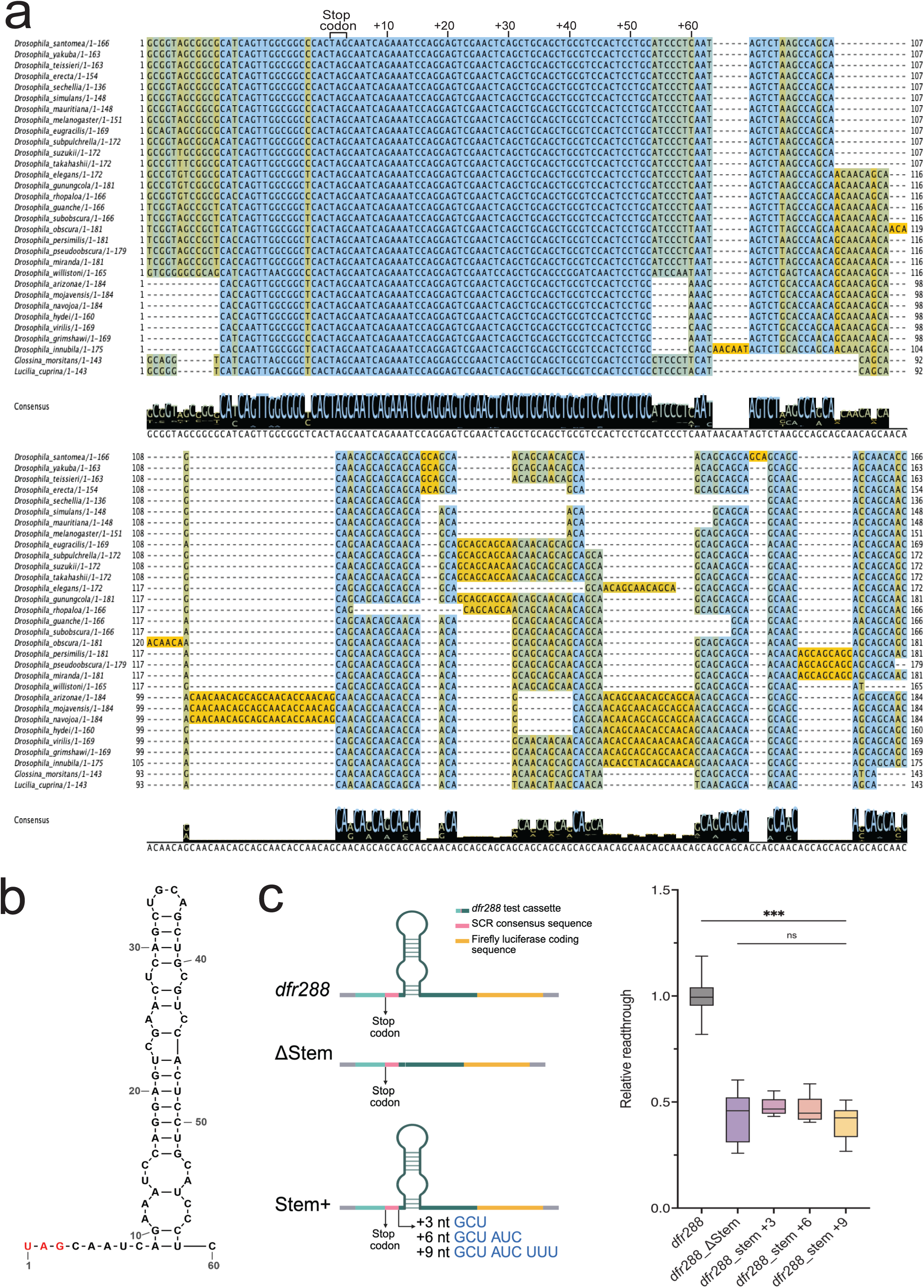
Stop codon identity and 3’ nucleotide context determine *dfr* translational readthrough. **a** Multiple sequence alignment depicting nucleotide conservation 30 nt upstream and 133 nt downstream of the first stop codon in the *dfr* gene. The blue-to-yellow gradient represents sequence conservation levels. The stop codon and the downstream 60 nt are indicated at the top of the aligned sequences. The guide tree for the alignment is available as supplementary material (Supp. Fig. S4). Illustration created with Jalview **b** Schematic representation of the *dfr 3’* mRNA stem-loop structure, predicted with RNAfold and mxFold2 WebServers, and redrawn using RNAcanvas. **c left** Schematic representation of the constructs analyzed in the readthrough assay, **right** Box plot diagram depicting relative readthrough efficiency observed with the indicated *dfr288*-based reporter constructs*: dfr288_Δstem* (deletion of the predicted stem-loop), *dfr288_stem+3* (3 nt inserted after +9 nt)*, dfr288_stem+6* (6 nt inserted after +9 nt)*, and* dfr288_stem+9 (9 nt inserted after +9 nt). In all box plot diagrams, boxes represent the 25-75% quartile, lines represent the median, whiskers represent min to max. Statistical significance was tested by one-way ANOVA with Dunnett’s T3 multiple comparisons test (*n*= 3 independent experimental replicates, \**\*\* = p* ≤ *0.001, n.s. = p* ≥ *0.05)*.

Several studies in various organisms have proposed that mRNA secondary structures formed by the mRNA’s 3’ UTR can enhance translational frameshifting and, in a few cases, also translational readthrough ^11, 24^. Intriguingly, our initial analysis uncovered an SCR-promoting effect of the *dfr* 3’ UTR sequence in the +39 to +189 region (Fig. 1c). To examine whether structural elements are present in the *dfr* 3’ UTR, we used the web-based prediction tools RNAfold and mxfold2 (Supp. Data SD2) ^32, 33^. Both tools predicted a thermodynamically stable stem-loop structure formed by the +9 to +59 nt sequence (Fig. 2b, Supp. Data SD2), confirming a previous prediction by the RNAz algorithm ^10^. As shown above, the predicted stem-loop sequence is completely conserved up to nt +53 in all the 29 *Drosophila* species analyzed here and in the two distantly related dipterans, strongly indicating functional conservation.

To experimentally test whether the predicted stem-loop structure plays a role in *dfr* SCR regulation, we analyzed a series of deletion and insertion constructs, designed to assess the presence and functionality of the SCR-promoting element, as well as the significance of its exact positioning. A specific deletion of the predicted stem-loop-forming nucleotide sequence (+10 to +59nt, *dfr288_*Δ*stem*), caused a significant decrease in SCR (Fig. 2c). When the predicted stem-loop was shifted further downstream by introducing either 3 nt, 6 nt, or 9 nt after the +9 nucleotide (*dfr288_stem+3; +6 or +9*)(Fig. 2c, left), we also observed a significant decrease in SCR (Fig. 2c, right). Notably, mRNA structure predictions did not reveal significant alterations of the 3’ UTR stem-loop in the stem+3/6/9 nt variants (Supp. Fig. S3, Supp. Table ST2) and none of these constructs exhibited decreased mRNA stability (Supp. Fig. S3). These findings confirm that a proximal stem-loop functions as an SCR enhancer and that its distance from the termination site critically influences readthrough efficiency.

### The thermodynamic stability of the mRNA secondary structure affects stop codon readthrough efficiency

To investigate the role of mRNA sequence identity *versus* mRNA secondary structure, we conducted site-directed mutagenesis to introduce nucleotide substitutions that alter the thermodynamic stability of RNA structures (Figure 3). We first designed three destabilizing variants (DV) by disrupting Watson-Crick base-pairing between opposing bases of the mRNA stem region (Fig. 3a and b; magenta arrowheads indicate mutations). Using RNAfold for structure predictions and free-energy calculations, we confirmed that all the destabilized stem-loop variants had an increase in Minimum Free Energy (MFE), indicating more dynamic and less stable structures (Supp. Table ST2). These DV variants showed significantly decreased levels of SCR efficiency (Fig. 3c). Importantly, the decrease was not caused by alterations in transcript levels (Fig. 3c, Supp. Fig. S3). Testing the same DV variants in a longer construct (*dfr453*), confirmed that the observed effects were not constrained by the length of the ORF (Supp. Fig. S3).

**Figure 3:**
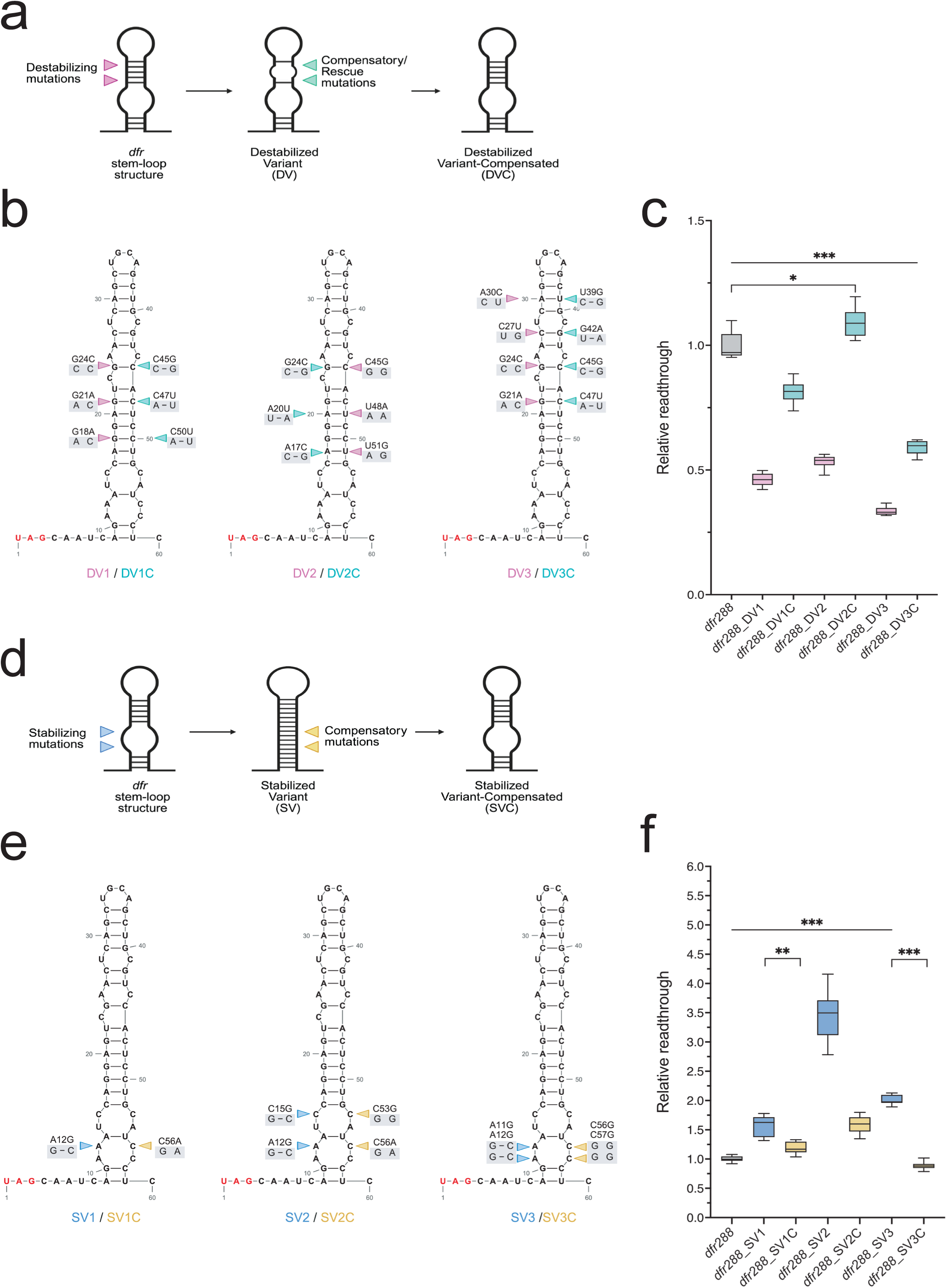
Altering the stem-loop stability affects *dfr* SCR efficiency. **a** Cartoon illustrating the mutagenesis strategy employed to destabilize the *dfr* mRNA stem-loop structure. Magenta arrowheads indicate destabilizing mutations, and cyan represents compensatory/rescue mutations. **b** Schematic depiction of the predicted *dfr* mRNA stem-loop structures. Magenta arrowheads indicate the mutations introduced to create the destabilizing variants; cyan arrowheads highlight the respective mutations introduced for compensatory variants. **c** Box plot diagram of the relative readthrough efficiencies (see methods for description) observed after transfection with the indicated *dfr_288*-based, destabilizing (DV) and structurally compensated (DVC) variants. **d** Cartoon illustrating the mutagenesis strategy used to stabilize the *dfr* mRNA stem-loop. Blue arrowheads show stabilizing mutations, and yellow indicates compensatory mutations. **e** Schematic depiction of the predicted *dfr* mRNA stem-loop structures. Blue arrowheads denote mutations for stabilizing variants; yellow arrowheads indicate mutations introduced to create compensatory variants that restore the initial *dfr288* stem-loop structure. **f** Box plot diagram of the Relative Readthrough Efficiencies observed after transfection with the indicated *dfr_288*-based, stabilizing (SV) and structurally compensated (SVC) variants. In all box plot diagrams, boxes represent the 25-75% quartile, lines represent the median, whiskers represent min to max. Statistical significance was confirmed by one-way ANOVA with Dunnett’s T3 multiple comparisons test (*n* = 3 independent experimental replicates, *\*\*\* = p* ≤ *0.001, ** = p = 0.002, * = p = 0.03*). Schematics were created with BioRender.com, RNA structures were redrawn using RNAcanvas.

We then restored the mRNA stem-loop structure by introducing compensatory nucleotide substitutions that recreate Watson-Crick base-pairing in the predicted RNA stem (Fig. 3a and b; DV1C, DV2C, DV3C, indicated by cyan arrowheads). These compensated variants showed partial or complete rescue of SCR efficiency (Fig. 3c), while transcript levels were not affected (Supp. Fig. S3). These findings underscore the importance of the mRNA secondary structure for *dfr* SCR, not the nucleotide sequence of the RNA stem, as it diverged even more in the compensated variants, arguing against the nucleotide sequence *per se* being an important determinant of readthrough levels.

Next, we took the opposite approach and introduced additional base-pairing in bulge sections of the stem-loop to increase the stability of the mRNA structure (Fig. 3d and e, mutations marked by blue arrowheads). Structure predictions and free-energy calculations of the three stabilizing variants (SV1-3) showed a decrease in MFE, indicating less dynamic and more stable structures (Supp. Table ST2). Compared to *dfr288*, all SVs resulted in 2- to 3-fold higher readthrough values (Fig. 3f). RT-qPCR analysis confirmed that the effect was not due to changed mRNA levels, but to an increase in translation efficiency over the stop codon (Supp. Fig. S3).

We then introduced compensatory mutations in RNA stem sections of the SVs, to reverse the stabilizing effect and to recreate the structure to its original form (SVC) (Fig. 3d and e, marked by yellow arrowheads) which led to significantly decreased readthrough levels (Fig. 3f). The observed decrease was not due to reduction in the stability of the reporter transcript (Supp. Fig. S3). These results strongly support that the thermodynamic stability of the RNA stem is more important than the exact mRNA sequence to stimulate *dfr* translational readthrough. Taken together, the structure-sequence analysis of the *dfr* 3’ UTR region demonstrates the existence of an mRNA stem-loop structure, and that its position and thermodynamic stability are direct determinants of SCR efficiency.

### The mRNA stem-loop in the proximal 3’ UTR of *dfr* promotes SCR *in vivo*

Next, we used the Gal4/UAS-system for tissue-specific expression ^34^ and developed a *Drosophila* model to analyze the capacity of the *dfr* 3’ mRNA stem-loop to regulate SCR *in vivo* (Figure 4). The *dfr288::T2A*, *dfr288_*Δ*stem::T2A*, *dfr288_SV2::T2A,* and the *dfr288_TAG>CAG::T2A* inserts (Fig. 4a) were shuttled into *pJFRC8-40XUAS-IVS-mCD8::GFP* ^35^. The vector was slightly modified to facilitate in-frame expression of a membrane-localized mCD8a::Dfr288 fusion protein, which serves as ratio-metric translation control. In the case of SCR, cytoplasmic GFP will be translated from the same transcript (Fig. 4b). Transgene expression was induced in the larval prothoracic gland (PG) using the *spok-Gal4.1.45* driver line (*spok>*) because this tissue has been shown to exhibit *dfr* SCR in late larval stages ^18^. In agreement with this observation, we noticed strong expression of mCD8a as well as of cytosolic GFP in PG cells of 3^rd^ instar larvae carrying the minimal SCR element *dfr288 (spok-Gal>dfr288,* Fig. 4c, quantification in Fig. 4g). In contrast, *spok>dfr288_*Δ*stem* larvae exhibited strongly reduced GFP-expression in their mCD8-labelled PG-cells (Fig. 4d, quantification in Fig. 4g), confirming our assumption that the mRNA secondary structure promotes SCR. In agreement with our previous results, *spok> dfr288_SV2* larvae expressing a reporter with a stabilized version of the mRNA stem-loop (Fig. 4e), revealed significantly increased GFP signals compared to *dfr288* (Fig. 4g) verifying our observation that stabilization of the mRNA stem-loop increases SCR efficiency. As expected, *spok>dfr288_TAG>CAG* larvae that expressed a reporter in which the *dfr288* stop codon had been changed to a CAG sense codon (Fig. 4f) exhibited the highest GFP/mCD8a expression ratio in PG cells. However, we also noticed lowered overall transgene expression in these larvae (Supp. Fig. S4).

**Figure 4:**
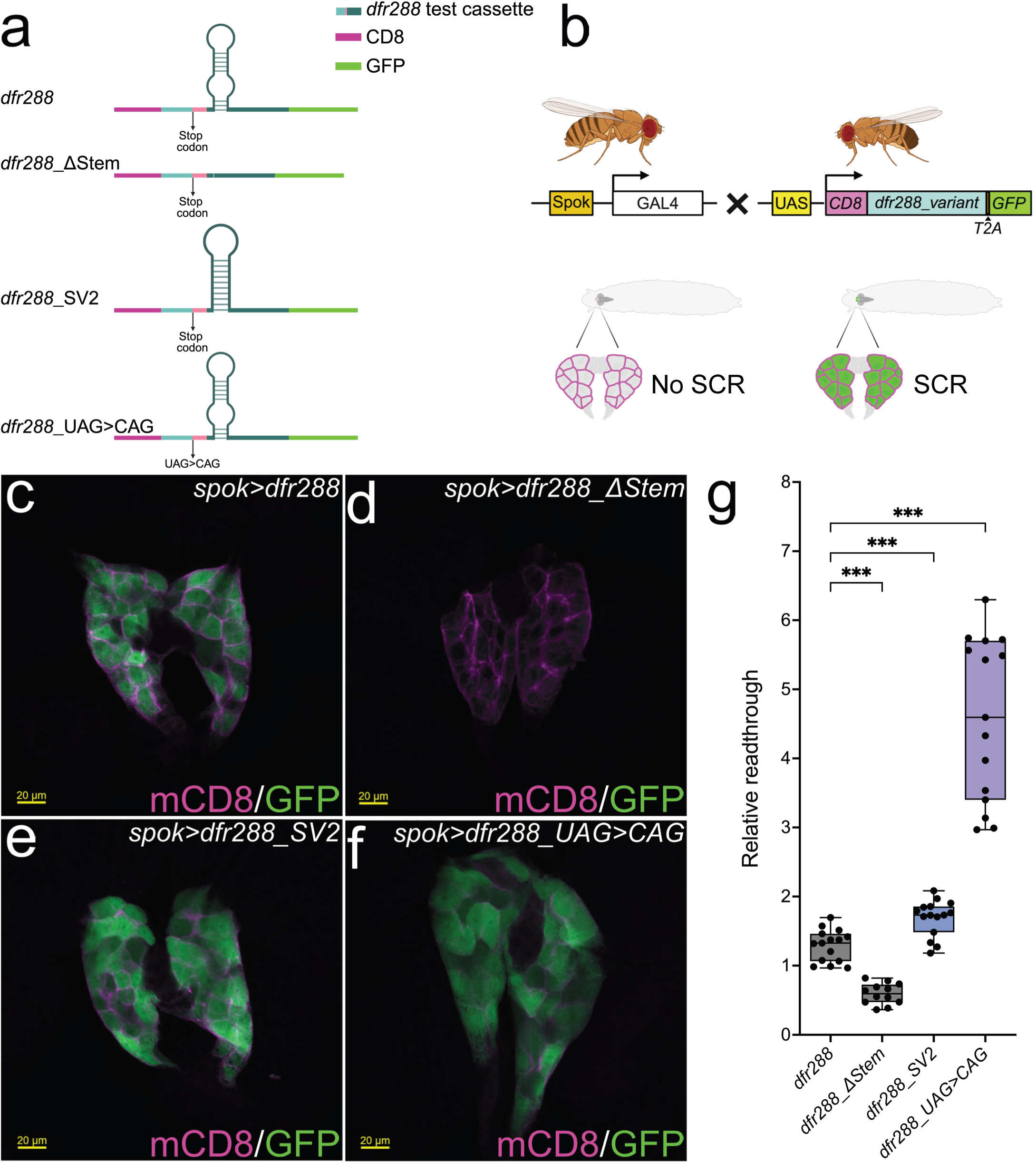
A 3’ UTR mRNA stem loop promotes *dfr* SCR *in viv*o. **a** Schematics of the *dfr288* reporter cassettes tested in the *in vivo* SCR assay. **b** Schematic representation of the Gal4xUAS system used for tissue-specific expression of the *dfr288* reporter transgenes. In F1 progenies of *spok-Gal4.1.45* (*spok>*) and flies carrying *UAS-dfr288* reporters (*UAS-CD8::dfr288_variant.T2A::GFP*), mCD8-expression is specifically induced in cells of the larval prothoracic gland (PG). Upon SCR, these cells co-express mCD8 and GFP. **c-f** Confocal images of anti-GFP (green) and anti-mCD8 (magenta) antibody staining of PGs from transgenic 3^rd^ instar larvae expressing *dfr288*-based reporters under the control of the *spok-Gal4.1.4.5* (*spok>*) driver. **c** *spok>mCD8::dfr288::T2A::GFP* exhibits PG-specific SCR revealed by GFP expression. **d** Reduced SCR in *spok>mCD8*::*dfr288_*Δ*stem::T2A::GFP* PG cells. **e** SCR in *spok>mCD8*::*dfr288_SV2::T2A::GFP* PGs. **f** SCR in *spok>mCD8*::*dfr288_TAG>CAG::T2A::GFP* larval PG cells. **g** Bee-swarm box and whiskers plot (min to max, all datapoints shown) of relative readthrough rates (represented as (RFI_GFP_/µm2)/(RFI_mCD8_/µm2)*ØRFI_mCD8_dfr288_/µm2) from max. intensity projections of identical volume confocal stacks from 3^rd^ instar larval PGs of the indicated genotype. Statistical significance was confirmed by Brown-Forsythe and Welsch ANOVA tests with Dunnett’s T3 multiple comparisons test (*n* = 12-15 larvae per genotype, *\*\*\* = p* ≤ *0.001*). Schematics were created with BioRender.com.

Although we cannot explain this effect, it could be speculated that the increase in translon length or the higher occurrence of T2A-containing polycistronic transcripts might have caused this negative effect on transgene expression. In conclusion, the presence of an mRNA stem-loop 3’ of the UAG stop codon significantly increased the SCR efficiency *in vivo*, and a more stable stem improved readthrough rates even further.

### Predictions of mRNA stem-loops in the 3’ UTR of genes with stop codon readthrough

To make a catalog of stop codons that may be affected by stem-loop structures, we used the structure-coding element in *dfr* 3’ UTR located between + 10 to + 58 as a guide and excised 60 nucleotides downstream of the first annotated stop codon of each *D. melanogaster* protein-coding gene in the reference genome (Fig. 5a). We then used computational structure prediction to find the most likely secondary mRNA structure of each of these sequences and selected the ones that confirm to a stringent definition of a stem-loop (Methods). In total, 1012 genes fulfilled these criteria and are candidates for stem-loop-facilitated SCR (Fig. 5b). Out of these, 34 genes overlapped with genes that have conserved ORFs downstream of stop codons ^10^ and 21 genes overlapped with genes that show evidence of SCR according to ribosome profiling by sequencing ^12^. In conclusion, we identified 50 genes (Fig. 5b and Supp. Table ST3) with predicted stem-loop-facilitated SCR in the +60 3’ UTR, including *dfr*, and out of these, 41 had not been predicted previously to contain mRNA stem-loops in the 3’ UTR (Supp. Table ST3).

**Figure 5:**
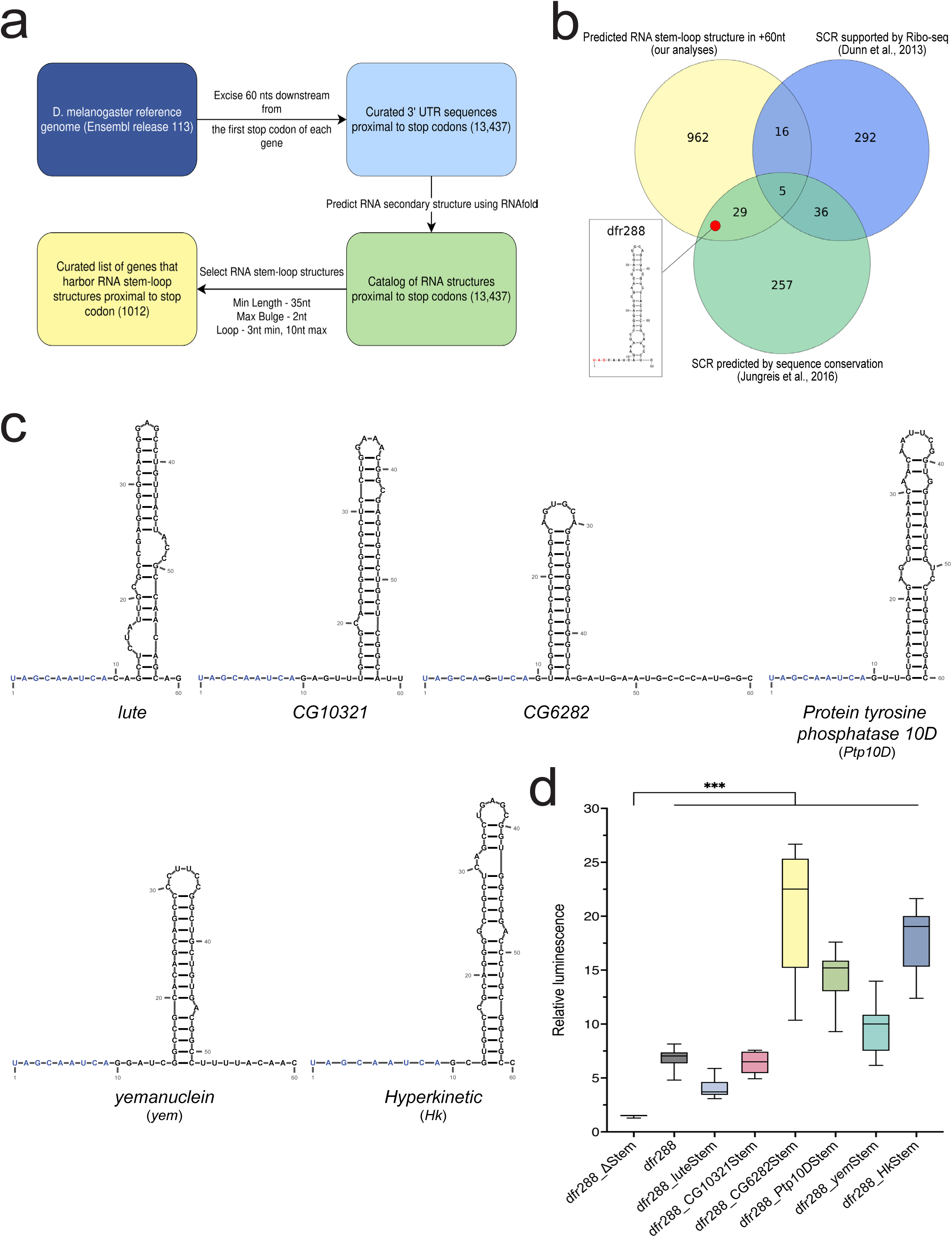
Prevalence and relevance of mRNA-secondary structures in *Drosophila melanogaster*. **a** Flowchart of our genome-wide computational screening for mRNA stem-loops downstream of stop codons. We considered only one sequence for each gene (mRNA). The minimum length of the stem-loop should be 35 nt; the terminal loop should be between 3 and 10 nt and no bulges (unpaired nucleotides within the stem) should be larger than 2 nt. **b** Venn diagram depicting sets of genes (mRNAs) with a predicted RNA stem-loop structure downstream of a stop codon (yellow circle), with evidence for SCR obtained from Ribo-Seq data of *Drosophila* embryos (blue circle), and with conserved ORFs downstream of stop codons (green circle). The *dfr288* stem-loop (box) is found in the yellow/green intersection because *dfr* SCR is not observed in embryos. **c** RNA-secondary structure prediction of the six mRNAs tested in the reporter assay. The +1 to +9 nt sequence color-coded in blue represents the *dfr* SCR consensus motif. **d** Box plot diagram of Relative Luminescence (FLuc/RLuc) values (see methods) of *Drosophila* S2 cells transfected with the six mRNA variants indicated on the X-axis. Boxes represent the 25-75% quartile, lines represent the median, whiskers represent min to max. Statistical significance was confirmed by one-way ANOVA with Dunnett’s T3 multiple comparisons test (*n* = 3 independent experimental replicates, *\*\*\* = p* ≤ *0.001*). Schematics were created with BioRender.com, RNA structures were redrawn using RNAcanvas.

To also analyze a somewhat longer 3’ UTR sequence we repeated the analysis with +80 nucleotides from each *D. melanogaster* protein-coding gene in the reference genome. As expected, the total number of genes that fulfilled the selection criteria increased to 1425 (Supp. Fig. S5), and 48 have been predicted to confer SCR by phylogenetic conservation and 31 by ribosome profiling. Thus, we identified 72 genes with a predicted stem-loop in the +80 region that overlaps with predicted or reported SCR, including *dfr* here as well (Supp. Fig. S5). As expected, there is a large overlap of genes identified in the +60 and +80 data sets (Supp. Fig. S5), and in total 78 genes were predicted to contain mRNA stem loops and been identified as SCR genes previously (Supp. Table ST3), and out of these, 68 had not been predicted previously to contain mRNA secondary structures in the 3’ UTR. Surprisingly, six genes were identified with 3’ stem-loops in the +60 but not in the +80 data sets. Further manual analyses of these six 3’ sequences revealed that at least two alternative stem-loops are predicted when +80 nucleotides are included. The ones selected by the RNAfold algorithm, based on MFE values, did not adhere to the selection criteria and was filtered away. Further experimental structure-function analysis of the predicted stem-loop structures will be needed for a true validation of their capacity to regulate SCR.

### Predicted mRNA stem-loops stimulate stop codon readthrough in *Drosophila* cells

To investigate the hypothesis that mRNA stem-loops in general can stimulate SCR, we experimentally analyzed six predicted stem-loops out of the 44 transcripts that had been identified through our genome-wide analysis in both the +60 and +80 data sets (Fig. 5b, Supp. Fig. S5). These six stem-loops are present in genes previously predicted to confer SCR by evolutionary conservation of protein coding sequences in the 3’ UTR ^10^. In addition, the gene for the Protein tyrosine phosphatase 10D (Ptp10D) has previously been show to confer programmed SCR in the *Drosophila* brain ^15^. We then inserted the predicted stem-loop forming sequences of these candidate genes in the *dfr288_*Δ*stem* construct while retaining the *dfr* minimal SCR-element (UAG stop codon and +4 to +9 nt sequence). This approach was intended to ensure that any observed changes in SCR are not due to alterations of the stop codon and its proximal sequence context but solely reflect the SCR-promoting capacity of the stem-loop. To account for potential structural changes introduced by modifying the +1 to +9 nt sequence of the tested genes, the secondary structures of each variant were reanalyzed using RNAfold (Fig. 5c, Supp. Data SD2). All six predicted stem-loops significantly enhanced SCR compared to the *dfr288_*Δstem variant (Fig. 5d). Notably, the stem-loops from *CG6282* and *Hk* exhibited SCR levels 4-5 times higher than that of *dfr288* itself, confirming that stem-loop structures with low MFE (Fig. 5d, Supp. Table ST2) enhance readthrough rate more efficiently. These results further support the hypothesis that mRNA stem-loop structures located in the 3’ vicinity of a stop codon, regardless of their genomic origin, can act as SCR-promoting elements.

## Discussion

Several modulatory mechanisms that can influence translational accuracy have been described including changes in cellular levels and structure of eukaryotic release factors, differences in ribosome complex composition, variations in the structure, abundance or identity of tRNAs, as well as post-transcriptional modifications of mRNA, rRNA, or tRNA ^6, 36, 37^. In certain cases, the identity of the nascent peptide has been found to influence the efficiency of translation termination ^38, 39^.These mechanisms influence the efficiency of stop codon recognition and translation termination fidelity, and might therefore explain general translational readthrough rates. Programmed SCR however, remains a relatively understudied gene regulatory process. It is hypothesized that programmed SCR of a specific mRNA requires combined regulation by *cis-* regulatory elements and *trans-*acting factors. Features such as the identity of the stop codon, the sequence context surrounding the stop codon, the presence of mRNA structural elements, and the interaction of mRNA-binding proteins have all been reported to contribute to SCR regulation ^24, 40-45^. However, a direct correlation between 3’ UTR mRNA architecture and its regulatory effect on SCR *in vivo* has not been demonstrated previously.

To study the molecular mechanisms underlying programmed SCR, we first mapped *cis-*regulatory elements in the mRNA encoded by *dfr,* a gene that exhibits spatio-temporally regulated SCR *in vivo* ^18^. In coherence with previous findings in other model systems, our data reveal that stop codon identity also acts as deterministic parameter of programmed SCR ^41^ with UGA providing the highest SCR rates, followed by UAG, and UAA which is the most efficient termination codon. This evolutionarily conserved hierarchy of termination codon efficiency possibly reflects inherent structural and chemical properties of the ribosomal machinery and/or the translation termination complex.

Interestingly, the +4 to +9 nucleotide sequence context downstream of the *dfr* stop codon matches the reported consensus found in polycistronic transcripts of certain plant viruses ^24, 27^, and our reporter assay indicates that this sequence is required but not sufficient for *dfr* SCR regulation. The +4 to +9 nucleotides are embedded in the ribosome’s mRNA entry channel when the stop codon is positioned in the decoding A site, and it has been shown that the identity of these nucleotides affect eukaryotic translation termination efficiency ^26, 45^. Thus, the high degree of sequence identity within the stop codon context most likely reflects the deep conservation of eukaryotic ribosomes and the mechanistic requirements for stop codon suppression to occur.

Multiple sequences alignment (MSA) constitutes an excellent tool for the identification of gene regulatory elements, based on the evolutionary conservation of DNA/RNA sequences in genomes that have diverged extensively from their common ancestor. Here we took advantage of existing whole genome sequences of 31 dipteran species with evolutionary distances spanning 40-60 Myr, and with an estimated substitution rate of 4.13 per neutral site ^46^. The MSA over the 1^st^ UAG stop codon and proximal 3’ UTR of *dfr* mRNA shows near 100% sequence identity over the identified mRNA stem-loop structure. This is in stark contrast to the nucleotide sequence further downstream, which also is highly conserved, but at the amino acid coding level, with frequent synonymous substitutions every 3^rd^ base (Fig. 2a). This strongly argues for functional importance of the -10 to +60 region covering the mRNA stem-loop, and that this RNA secondary structure constitutes an evolutionarily ancient regulatory element of *dfr* mRNA translation.

RNA secondary structures, especially stem-loops, have been shown previously to affect genetic re-coding events like ribosome frameshifting, selenocysteine incorporation, and programmed SCR ^24, 47^. Earlier studies on *Drosophila hdc* and *kel* employed deletion mapping and analysis of heterologous mRNA to confirm that the predicted, stem-loop containing regions impact SCR *in vitro* and *in vivo* ^13, 15^; however, this does not clarify if a stem-loop structure or a specific sequence motif is required. We therefore thoroughly investigated the relative contribution of mRNA secondary structure versus primary sequence context in *Dfr* SCR regulation using mutational analysis. Reducing the base pairing within the *dfr* 3’ stem-loop distinctly lowered SCR efficiency while closing bulges markedly increased readthrough rates. Importantly, compensatory mutations could specifically reverse these effects although additional changes in the primary sequence context were introduced.

Finally, replacing the *Dfr* SCR promoting element with non-related, stem-loop forming sequences obtained from our *in silico* prediction maintained or even increased SCR of the heterologous mRNAs, pin-pointing the key importance of mRNA stem-loop structures in determining SCR efficiency. While this is the first report highlighting a direct and positive correlation between mRNA secondary structure stability and the efficacy of translational readthrough, the molecular mechanism by which mRNA stem-loops promote programmed SCR remains elusive. It has been suggested that mRNA secondary structures prolong ribosome dwell time, thereby increasing the chance of non-standard decoding events such as frameshifting and codon re-assignment ^48^. It is conceivable that ribosomal dwell time would be increased in the presence of mRNA secondary structures with higher thermodynamic stability because such structures would assemble faster, require more energy to be resolved, and are more likely to re-assemble between individual rounds of translation. Interestingly, recent cryo-EM data reveal direct interactions between the ribosome and the SARS-CoV-2-RNA-pseudoknot, which increases ribosomal dwell time and stimulates frameshifting ^49^, suggesting that a similar scenario could exist during programmed SCR.

Overall, our data reveal that regulation of *dfr* SCR is a complex interplay between stem-loop stability, distance of the stem-loop from the stop codon, stop codon identity, and +4 to +9 sequence context. From an evolutionary perspective, gene-specific fine-tuning of programmed SCR by various elements would be favored because the prevalent, canonical gene product usually has essential roles during development, which might be compromised by precocious or overabundant SCR. In addition, the SCR gene product might only be required at a specific time-point and under certain conditions. This seems to be the case for Dfr-L which has been functionally associated with steroid hormone metabolism and immunity ^18^. Finally, variability in mRNA stem-loop geometry and position would allow interactions with a range of trans-acting factors providing another layer of spatio-temporal SCR regulation.

Phylogenetic analyses to discover evolutionarily conserved mRNA secondary structures involved in SCR regulation are frequently conducted using the RNAz algorithm ^50^. In our refined approach, we combined RNAfold in *D. melanogaster* and a more stringent set of structural parameters obtained from our *dfr* SCR characterization as filtering criteria, which identified 68 new candidates for stem-loop-facilitated SCR (Supp. Table ST3). Together with the 24 genes previously predicted to confer SCR and also predicted to have mRNA structures in the 3’ UTR ^10^ this sums up to 92 genes in *D. melanogaster* prone to SCR that also contain 3’ mRNA stem-loop structures.

Our experimental validation of six of these new mRNA stem-loops conferred their SCR-stimulatory capacity in a cell-based assay system. This clearly demonstrates that mRNA stem-loops in the proximal 3’ UTR can promote SCR of a heterologous gene construct, and we conclude that mRNA stem loops may constitute a more general mechanisms for SCR regulation than previously anticipated.

While SCR luciferase reporter assays are an efficient method to analyze large numbers of sequence variants, they cannot provide the complete picture of programmed SCR regulation in multicellular organisms. We therefore confirmed the SCR-promoting capacity of the stem-loop structure in a transgenic *Drosophila* model, which recapitulated the results from the transfected S2 cells. Comparing SCR efficiencies in PG cells reveals about 30% SCR upon *spok>dfr288* expression in relation to the *spok>dfr288_UAG>CAG* sense codon control. This readthrough efficiency is substantially higher than the ∼18% relative SCR efficiency from the same sequences analyzed in cultured S2 cells (Fig. 1f). While we would like to suggest that this could reflect the activity of tissue-specific regulatory factors in the PG cells, we cannot exclude the possibility that reporter gene activities might differ between our two-plasmid luciferase system and the single-unit mCD8/GFP *in vivo* reporter. Similarly, concerns regarding the use of dual reporter systems that rely on independent constructs, or fused reporter proteins have been raised ^25, 51^, which strongly recommends the use of complementary reporter systems to minimize experimental artifacts. The high rate of *dfr* SCR in PG cells correlates well however, with previously reported readthrough levels of the endogenous *dfr* gene ^18^. In that study, western blot analysis showed that expression of the readthrough product Dfr-L (derived from translon 2) in PG cells is about 50% in comparison to the canonically translated Dfr-S (derived from translon 1), which would equal to about 33% of Dfr-L in relation to the total amount of Dfr protein expressed in PGs. Thus, the transgenic *dfr288* construct confers readthrough levels that are in the same range as the endogenous expression level of Dfr-L *in vivo*, suggesting that the *dfr288* construct contains all the regulatory *cis-*acting elements required to promote efficient SCR of *dfr* in PG cells.

The high level of gene-specific SCR of natural stop codons as described here for *dfr* SCR, with *in vivo* readthrough rates greater than 30%, could guide attempts to enhance readthrough efficiency for biotechnological and pharmacological purposes. Further mechanistic understanding and cross-species evaluation is needed however, before such approaches would be broadly applicable.

## RESOURCE AVAILABILITY

### Lead Contact

Further information and requests for resources and reagents should be directed to and will be fulfilled by the lead contact, Ylva Engström (mail to).

### Materials Availability

Fly lines and plasmids generated in this study are available upon request from the lead contact.

### Data and code availability

- The script for mRNA stem-loop prediction is publicly accessible on GitHub (https://github.com/iealexop/detect_RNA_hairpin), and on Zenodo (doi:10.5281/zenodo.16782268).
- All original data reported in this paper will be shared by the lead contact upon request.
- Any additional information required to analyze the data reported in this paper is available from the lead contact upon request.

## Author contributions

Conceptualization and designing the study: L.K., G.W., M.R.F. and Y.E.; Methodology, Investigation, and Formal analysis: L.K., G.W., D.W.; Software and Data Curation: I.A. and A.E.; Writing-Original draft: L.K., G.W., M.R.F. and Y.E.; Writing-Review & Editing: L.K., G.W., D.W., I.A., A.E., M.R.F. and Y.E.; Supervision: M.R.F. and Y.E.; Funding Acquisition: M.R.F and Y.E.

## Acknowledgement

The authors would like to thank Kim Rewitz for generously sharing the *spok-Gal4.1.45* driver. Experimental resources obtained from the Bloomington *Drosophila* Stock Center (NIH P40OD018537) and the *Drosophila* Genomics Resource Center (NIH Grant 2P40OD010949) were used in this study. The computations were enabled by resources provided by the National Academic Infrastructure for Supercomputing in Sweden (NAISS), partially funded by the Swedish Research Council through grant agreement no. 2022-06725. The authors also acknowledge the technical support from the IFSU (Imaging Facility at Stockholm University). The authors would like to thank the FlyBase consortium for curating and providing crucial information about *Drosophila* genetics and experimental tools.

## Material and Methods

### Plasmid construction

To generate *pAct5c-MCS-T2A-Fluc,* the *pAct5C-MCS* ^52^ backbone, the *Firefly luciferase (FLuc)* sequence from *pJD375* ^53^, and the T2A sequence from *pPGxRF3* (Addgene #138384) were used. The *FLuc* start codon was mutated using site-directed mutagenesis by PCR ^54^. A list of all primers used in this study is available as Supplementary Table ST4. The Phusion™ High-Fidelity DNA Polymerase (ThermoFisher, #F530S) was used to amplify fragments of the *dfr locus* from *w^1118^* genomic DNA. PCR-products were added to the *pAct5c-MCS-T2A-FLuc* reporter by conventional restriction/insertion cloning. To generate the *dfr288* variants shown in Figures 2 and 3, PCR-directed site-directed mutagenesis was used ^54^. The *dfr288* stem swap variants with stem-loop structures from *lute, Ptp10D, Hk, yemanuclein, CG10321* and *CG6282* genes were made with the Q5®-site directed mutagenesis kit (NEB, #E0554S) according to the manufacturer’s instructions. The *Copia Renilla* Control plasmid (Addgene #38093) was used to measure transfection efficiency. To allow in-frame expression of the dfr288::T2A inserts with mCD8a and GFP from *pJFRC8-40XUAS-IVS-mCD8::GFP* (Addgene #26221)^35^, we modified the vector using the primer pair *Q5SDM_pJFRC8_BamHIOOF_F*, and *Q5SDM_pJFRC8_BamHIOOF_R* in a Q5®-mutagenesis PCR reaction (NEB, #E0554S). The inserts *dfr288::T2A*, *dfr288_*Δ*stem::T2A*, *dfr288_TAG>CAG::T2A* and *dfr288_SV2::T2A* were shuttled between *pAct5c-MCS-T2A-Fluc* and the modified *pJFRC8-40XUAS-IVS-mCD8::GFP* plasmid using a *BamHI* restriction site located between the *CD8a-* and *GFP*-coding sequences. All constructs were verified by Sanger sequencing (Eurofins Genomics).

### Cell culture and transfection

Cell transfection and luciferase assays were performed in the *Drosophila* S2 cell line (DGRC Stock 6; https://dgrc.bio.indiana.edu//stock/6; RRID:CVCL_TZ72; ^55^). Cells were cultured at 25°C in Schneider’s Medium (Gibco, # 21720024), supplemented with 10% Fetal bovine serum (FBS), 50 units/ml penicillin, and 50 μg/ml streptomycin.

For luciferase assays, cells were seeded in a 96-well plate at a concentration of 1.0 x 10^4^ a day before transfection. After 1 day of incubation, transfection was carried out using the Effectene Transfection Reagent kit (QIAGEN, #301425), following the manufacturer’s instructions. Luciferase assays were carried out 48 hours after transfection.

### Luciferase assay

Luciferase assays were performed using the Dual-Glo® Luciferase Reporter Assay System (Promega, #E2920) following the manufacturer’s instructions. Luminescence was measured with an EnSpire 2300 Multi-mode microplate reader (PerkinElmer). The raw luminescence values were first corrected for background, and used for further calculations. *Relative Luminescence* values were obtained by normalizing Firefly luciferase (FLuc) luminescence values to the Renilla luciferase (RLuc) luminescence values (FLuc/RLuc). *Readthrough Efficiency* was calculated as the FLuc/RLuc values from the different test constructs relative to the FLuc/RLuc values of the *dfr288_UAG>CAG* sense codon control reporter. *Relative Readthrough Efficiency* refers to the readthrough efficiency gained from each construct in relation to the *dfr288 construct,* which was set to 1 in the box plots.

### dsRNA synthesis and RNAi in S2 cells

The templates for dsRNA synthesis of *Upf1* and *eRF1* were PCR-amplified from *w^1118^* genomic DNA, and for *RFP* from the pAWR Gateway vector. The dsRNA was synthesized using the T7 RiboMAX™ Large Scale RNA Production Systems kit (Promega #P1300) following the manufacturer’s protocol.

For the RNAi, S2 cells were seeded one day prior to addition of the dsRNA in a 96-well plate at a concentration of ∼3 x 10^4^ cells. The following day, the cells were starved in serum-free Schneider’s media for 3 hours. Subsequently, dsRNA was introduced to the cells in complete media at a concentration of 1 µg per well. After a 1-hour recovery period, the cells were transfected with the appropriate readthrough reporter constructs, and luminescence was measured 48 hours after initiation of the dsRNA treatment.

### RT-qPCR

For RT-qPCR assays, cells were seeded in a 48-well plate at a concentration of 3.0 x 10^4^. A day after seeding, the cells were transfected with the Effectene transfection reagent and incubated for 48 hours. RNA was extracted with TRIzol (Invitrogen, #15596018), cDNA synthesis was carried out using the SuperScript III Reverse Transcriptase (Invitrogen, #18080044) with Random Hexamers (ThermoScientific, #S0142), following the manufacturer’s protocol.

qPCR was performed in a Rotor-Gene Q system (QIAGEN) using Fast SYBR™ Green Master Mix (AppliedBiosystems, #4385614). The Cycle threshold (Ct) values of target genes in each run were determined using a standard curve generated by pooling cDNA from all the samples used in the analysis. The values obtained were normalized to *rpl32.* The comparative cycle threshold (2^−ΔΔ^CT) method was used to determine the fold change in *luciferase* reporter expression between transfected and untransfected samples.

### Whole-genome multiple alignments

Protein-coding sequences in FASTA format were downloaded from Ensembl Metazoa (https://metazoa.ensembl.org/index.html) for 29 *Drosophila* species and two outgroup flies (*Glossina morsitans* and *Lucilia cuprina*) (Supp. Table ST1). The longest protein isoform per gene was selected to identify conserved single-copy orthologs using BUSCO v5.4.7 (lineage: diptera_odb10) in protein mode, executed within a Docker environment (v28.0.4, build b8034c0) ^56^. The concatenated supermatrix of BUSCO-derived protein alignments was used for maximum likelihood phylogenetic inference with IQ-TREE2 ^57^, with the best-fitting model (Q.insect+F+R6) selected by ModelFinder. Whole-genome FASTA files were softmasked using RepeatMasker v4.1.6 prior to alignment, using species-specific repeat libraries where available. The softmasked genomes were then aligned using the Progressive Cactus pipeline (v2.9.8) ^58^.The final alignment was output in HAL format and processed using the HAL tools package ^59^ for downstream analyses, including liftover, slicing, and export to other formats.

### Fly husbandry and generation of transgenic fly lines

Flies were reared on potato-mash culture medium at 25°C, 55% relative humidity, and a 12:12 h day/night-cycle. Information about *Drosophila* genetics is available on the FlyBase Database of *Drosophila* Genes and Genomes ^60^. Transgenic fly lines were created by φ31-mediated, site-specific transformation ^61^. *pJFRC8-40XUAS-IVS-mCD8::dfr288::T2A::GFP*, *pJFRC8-40XUAS-IVS-mCD8::dfr288_*Δ*stem::T2A::GFP*, *pJFRC8-40XUAS-IVS-mCD8::dfr288_TAG>CAG::T2A::GFP* and *pJFRC8-40XUAS-IVS-mCD8::dfr288_SV2::T2A::GFP* plasmids were injected into *y^1^ w^1118^; PBac{y^+^-attP-9A}VK00022* flies (BDSC 9740) by BestGene Inc., and insertion events were identified by the presence of the *pJFRC8*-associated *w[mC+]* marker. Other *Drosophila* stocks used in this study: *w*; spok-Gal4.1.45* ^62^.

### Antibody-staining of *Drosophila* larvae

Whole-filet dissections of age-matched 3^rd^ instar *Drosophila* larvae (84 h +/-1 h after hatching) were fixed for 30 min in 4% formaldehyde in 1x phosphate-buffered saline (PBS) while gently rocking at room temperature (RT). Samples were briefly rinsed with 1x PBS supplemented with 0,2% Triton-X100 before rocking 10 min in 1xPBS+1% Triton-X100 at RT. After rinsing with 1x PBS+0,2% Triton-X100, samples were blocked for 30 min in 1x PBS+0,2% Triton-X100 supplemented with 5% normal goat serum (NGS, Invitrogen, #31873) at RT. Primary antibody incubation with anti-mouse CD8a Monoclonal Antibody 53-6.7 (eBioscience^TM^, purchased from Invitrogen #14-0081-82, and applied at a 1:200 dilution) and anti-GFP (Invitrogen, #A-11122, 1:500 dilution) in 1x PBS+0,2% Triton-X100+2,5% NGS was done overnight gently rocking at 4°C. As secondary antibodies, we used Anti-rabbit AlexaFluor488 and Anti-rat AlexaFluor594 (Thermofisher #A32790 and #A11007) in combination with DAPI (Sigma-Aldrich, #D9542; 2 mg/ml, applied at a 1:500 dilution). Samples were washed in 1x PBS+0,2% Triton-X100 for 3x 20 min gently rocking at RT before fine-dissecting and mounting in Fluoromount-G (Invitrogen, #00-4958-02). Stained samples were analyzed under a Zeiss LSM800 microscope with a Plan-Apochromat 20x/0.8 objective using ZEN Blue V2.1 software. Confocal stacks of 5 sections from prothoracic glands (PGs) were acquired and fluorescence intensities of the whole organ area were measured from maximum intensity projections. For quantification, mCD8 and GFP signal intensities were normalized to the measured area, and GFP/mCD8 ratios were calculated. Although we used the same expression vector for all constructs and chose the same genomic landing site, we noticed decreased mCD8 expression in *spok>dfr288_TAG>CAG* and to a lesser degree in *spok>dfr288_*Δ*stem* animals (Supp. Fig. S4). We therefore adjusted all sample GFP/mCD8 ratios to the average dfr288 mCD8 intensity level prior to statistical analysis with GraphPad Prism software (v10.5.0).

### mRNA stem-loop prediction

Reference *D. melanogaster* genome and annotation (BDGP6.46, Ensembl release 113) were retrieved from Ensembl and used to compile a FASTA file containing the most upstream 3’ UTR corresponding to each annotated gene. After trimming each sequence to maximum 60 nts length or 80 nts respectively, we used RNAfold (part of ViennaRNA package, version 2.4.17) to predict their secondary structure and print the output in dot-bracket notation, as ‘RNAfold -j -i first_3utrs_trimmed.fa > RNAfold_60.txt’. To determine whether these structures represent stem-loops, we used Python script that parses each dot-bracket notation and labels sequences as stem-loops if they have a minimum length of 35 nts; a loop region length ranging from 3 to 10 nts and bulges with maximum length of 2 nts. The script is publicly accessible on GitHub: https://github.com/iealexop/detect_RNA_hairpin and on Zenodo: doi:10.5281/zenodo.16782268

## Supplementary Figure Legends

**Supplementary Figure S1: a and b** Column graphs showing relative expression levels of the reporter transcripts in *Drosophila* S2 cells transfected with the indicated constructs (X-axis), as measured by RT-qPCR. Expression is normalized to *rpl32* mRNA and presented as fold change relative to untransfected cells. Data represent the mean± SEM from three independent biological replicates. Statistical significance was confirmed by one-way ANOVA with Dunnett’s T3 multiple comparisons test (*\* = p = 0.03,* ns *= p* ≥ *0.05)*.

**Supplementary Figure S2:** Evolutionary guide tree inferred from multiple sequence alignment of 29 Drosophila species and two additional out group flies based on their protein orthologs.

**Supplementary Figure S3: a** Schematic redrawing (BioRender.com) of mxFold2-predictions of mRNA secondary structures in the *dfr288*_Stem+3, +6, and +9 sequence variants. The predicted slight variation between the stem-loops results from the algorithm’s priority to select the structure with the lowest MFE. Because of the negligible differences in MFE between all predicted stem-loops (Supp Table ST2), it is more likely that identical stems are formed. **b-e** Column graphs depicting relative expression levels of the reporter transcripts in *Drosophila* S2 cells transfected with the indicated constructs (X-axis), as measured by RT-qPCR. Expression is normalized to *rpl32* mRNA and presented as fold change relative to untransfected cells. Data represent the mean± SEM from three independent biological replicates. Statistical significance was confirmed by one-way ANOVA with Dunnett’s T3 multiple comparisons test (*\*\* = p = 0.002, * = p = 0.03*, ns *= p* ≥ *0.05)*. **f, g** Box plot diagram of the Relative Readthrough Efficiencies (see methods) after transfection of *Drosophila* S2 cells with the respective *dfr_453*-based, destabilizing (DV) (**f**) and stabilizing (SV) (**g**) variants. In all box plot diagrams, boxes represent the 25-75% quartile, lines represent the median, whiskers represent min to max. Statistical significance was confirmed by one-way ANOVA with Dunnett’s T3 multiple comparisons test (*n* = 3 independent experimental replicates, *\*\*\* = p* ≤ *0.001, ** = p = 0.002, * = p = 0.03*).

**Supplementary Figure S4:** Column graph depicting the mean, area-normalized, relative fluorescence intensities (RFI/µm^2^) of mCD8 (pinks) and GFP (greens), revealed by immunofluorescence staining, in PGs of *Drosophila* L3 larvae expressing the indicated *dfr* SCR reporter transgenes under *spok-Gal4.1.45* driver control.

**Supplementary Figure S5: a** Venn diagram depicting sets of genes (mRNAs) with a predicted RNA stem-loop structure in the +80 nt region, downstream of a stop codon (yellow circle), with evidence for SCR obtained from Ribo-Seq data of *Drosophila* embryos (blue circle), and with conserved ORFs downstream of stop codons (green circle). **b** Euler diagram illustrating the intersection among gene sets (mRNAs) identified to have an RNA stem-loop structure within the 3’ UTR with two length parameters, +60 nt *vs* +80 nt. **c** Euler diagram illustrating the overlap between the predicted structures derived from +60 and +80 nt gene sets, which were also demonstrated or predicted to exhibit stop codon readthrough. **d** Column graphs depicting relative expression levels of the reporter transcripts in *Drosophila* S2 cells transfected with the indicated stem-loop variants (X-axis), as measured by RT-qPCR. Data represent the mean± SEM from three independent biological replicates. Statistical significance was confirmed by one-way ANOVA with Dunnett’s T3 multiple comparisons test (*\*\* = p = 0.002,* ns *= p* ≥ *0.05*).

## Additional files

Supplementary Table ST1-Information about sequences downloaded from Ensembl Metazoa for 29 *Drosophila* species and two outgroup flies (*Glossina morsitans* and *Lucilia cuprina*).

Supplementary Table ST2-Minimum free energy calculation of *dfr* mRNA secondary structures.

Supplementary Table ST3-List of genes predicted to form mRNA secondary structures in the genome-wide analysis of *Drosophila melanogaster*.

Supplementary Table ST4-List of primers used in this study.

Supplementary Data SD1-Multiple sequence alignment depicting nucleotide conservation of *dfr* ORF2. The blue-to-yellow gradient represents the levels of sequence conservation. The putative *cis-*acting elements identified are indicated on the top of the aligned sequences. Illustration created with Jalview and Biorender.

Supplementary Data SD2-Original predictions from RNAfold and mxfold2.

