## Supplementary figures and images for "The mRNA architecture of the termination site primes programmed stop codon readthrough events in *Drosophila*"

### Supplementary Data SD1

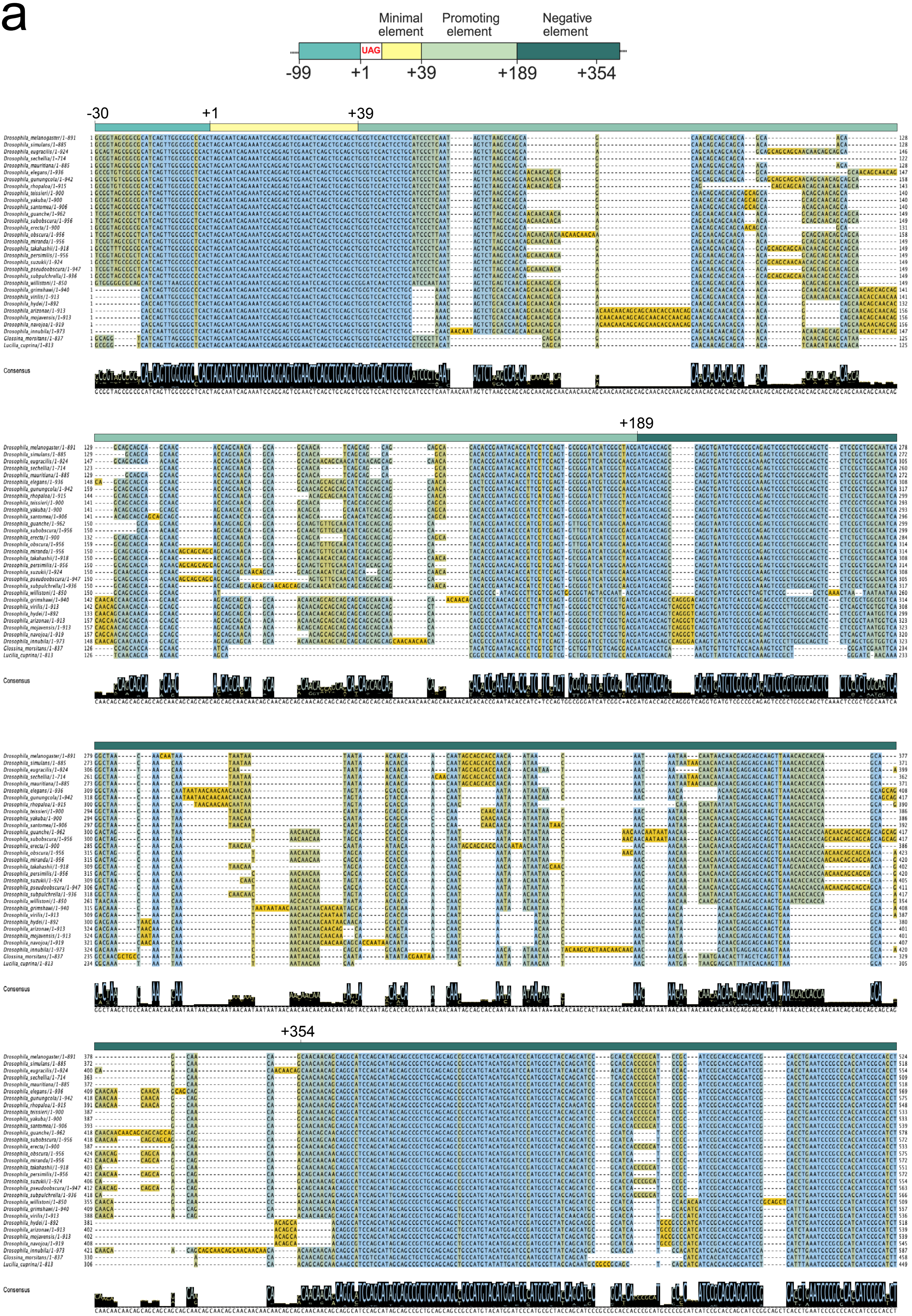

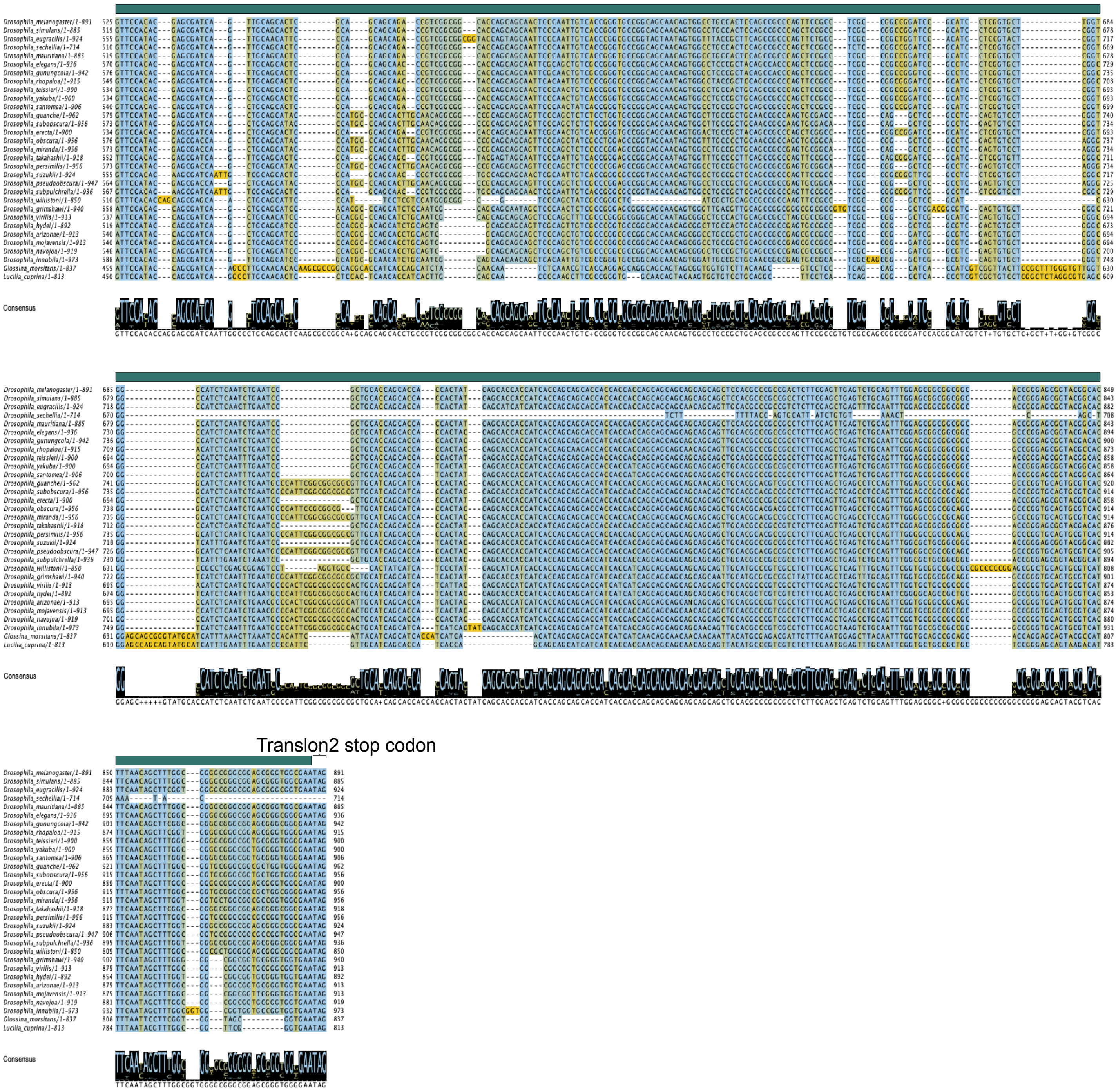

### Supplementary Figure S1

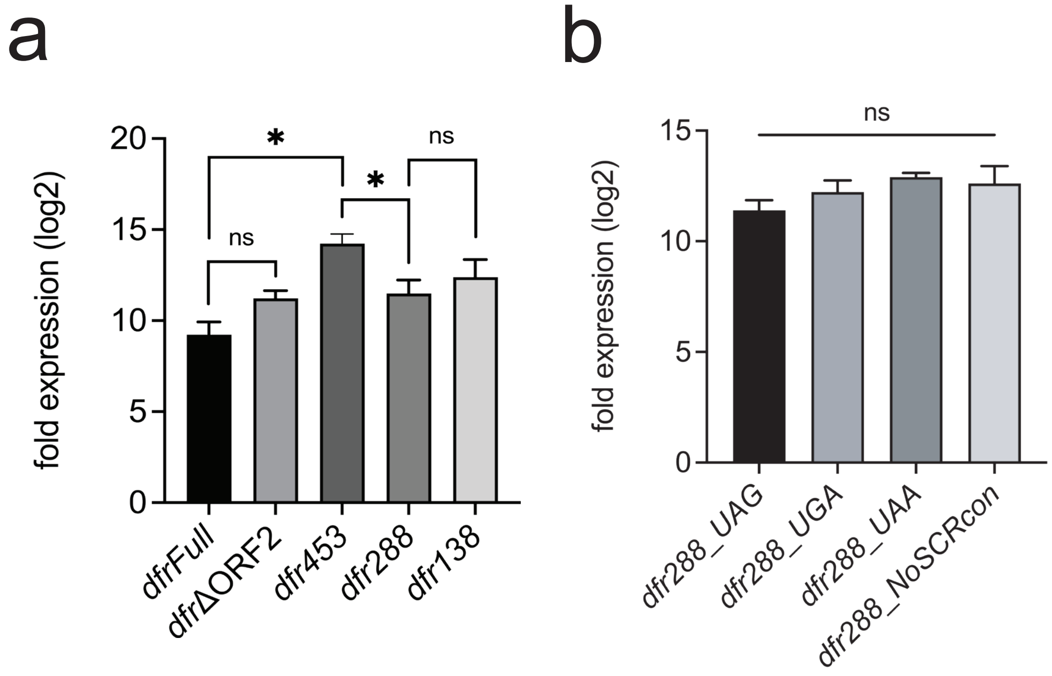


Supplementary Fig S1

### Supplementary Figure S2

a

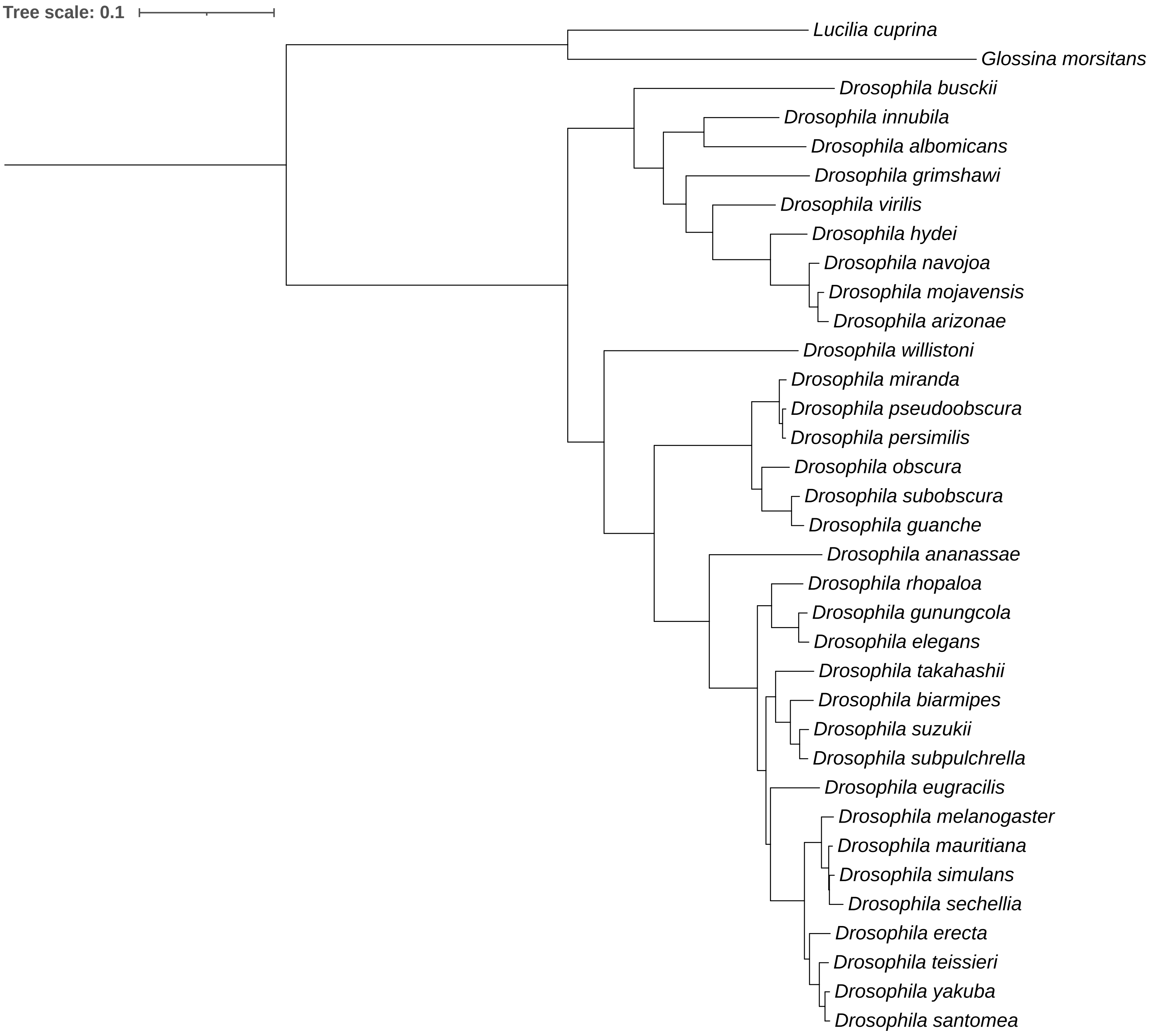


Supplementary Fig S2

### Supplementary Figure S3

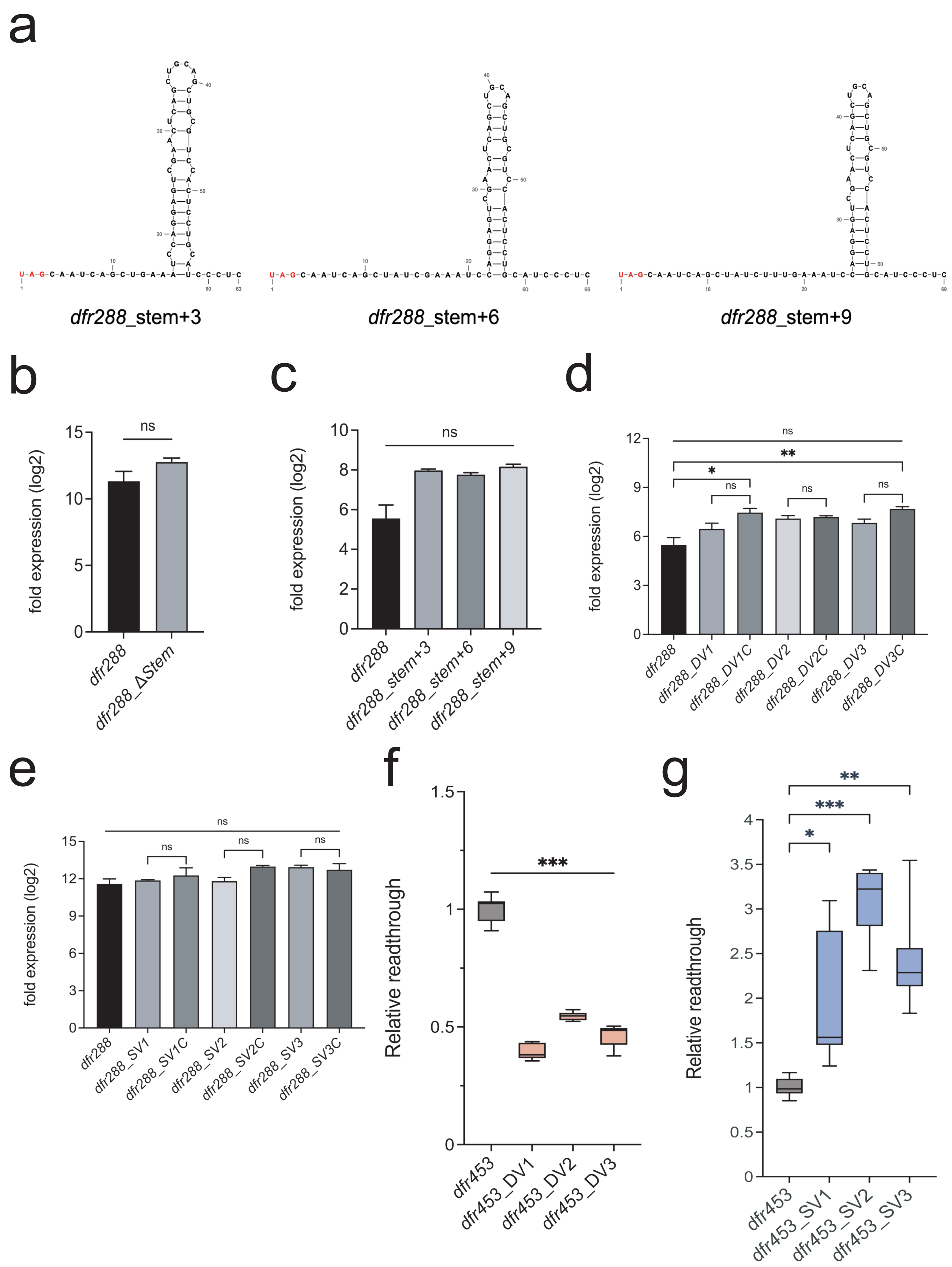


Supplementary Fig S3

### Supplementary Figure S4

a

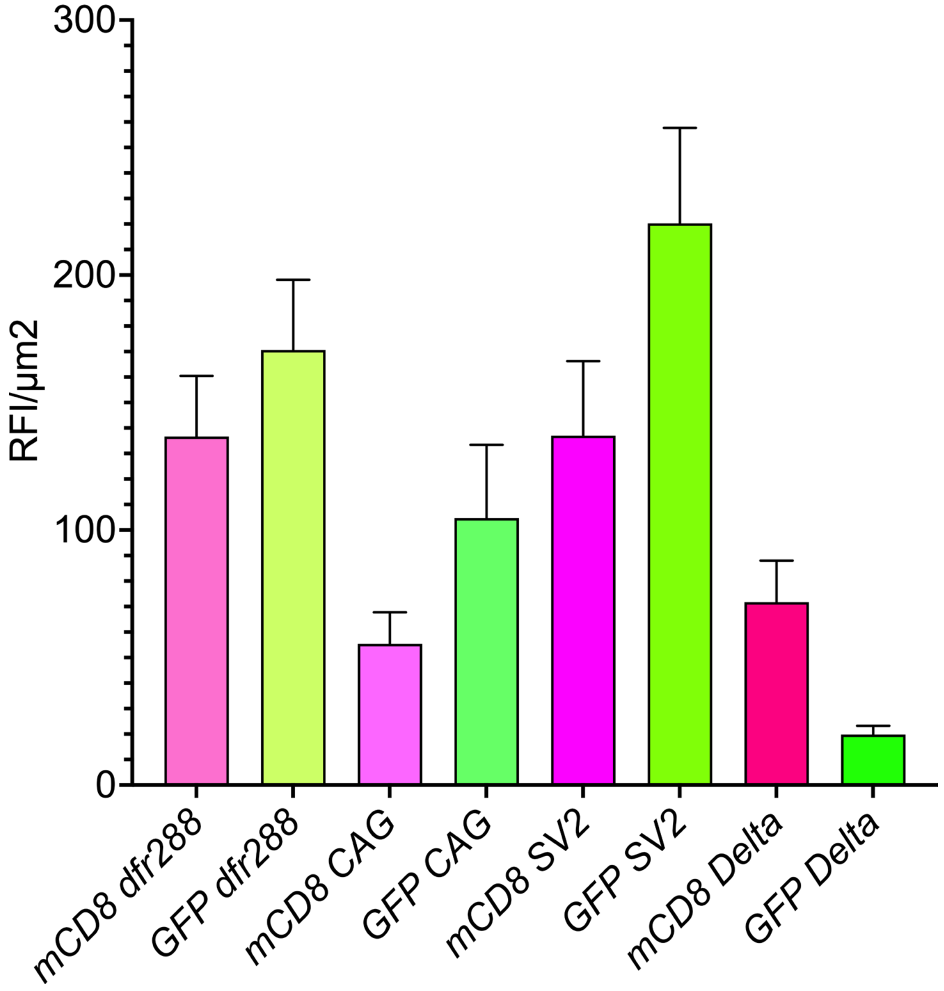


Supplementary Fig S4

### Supplementary Figure S5

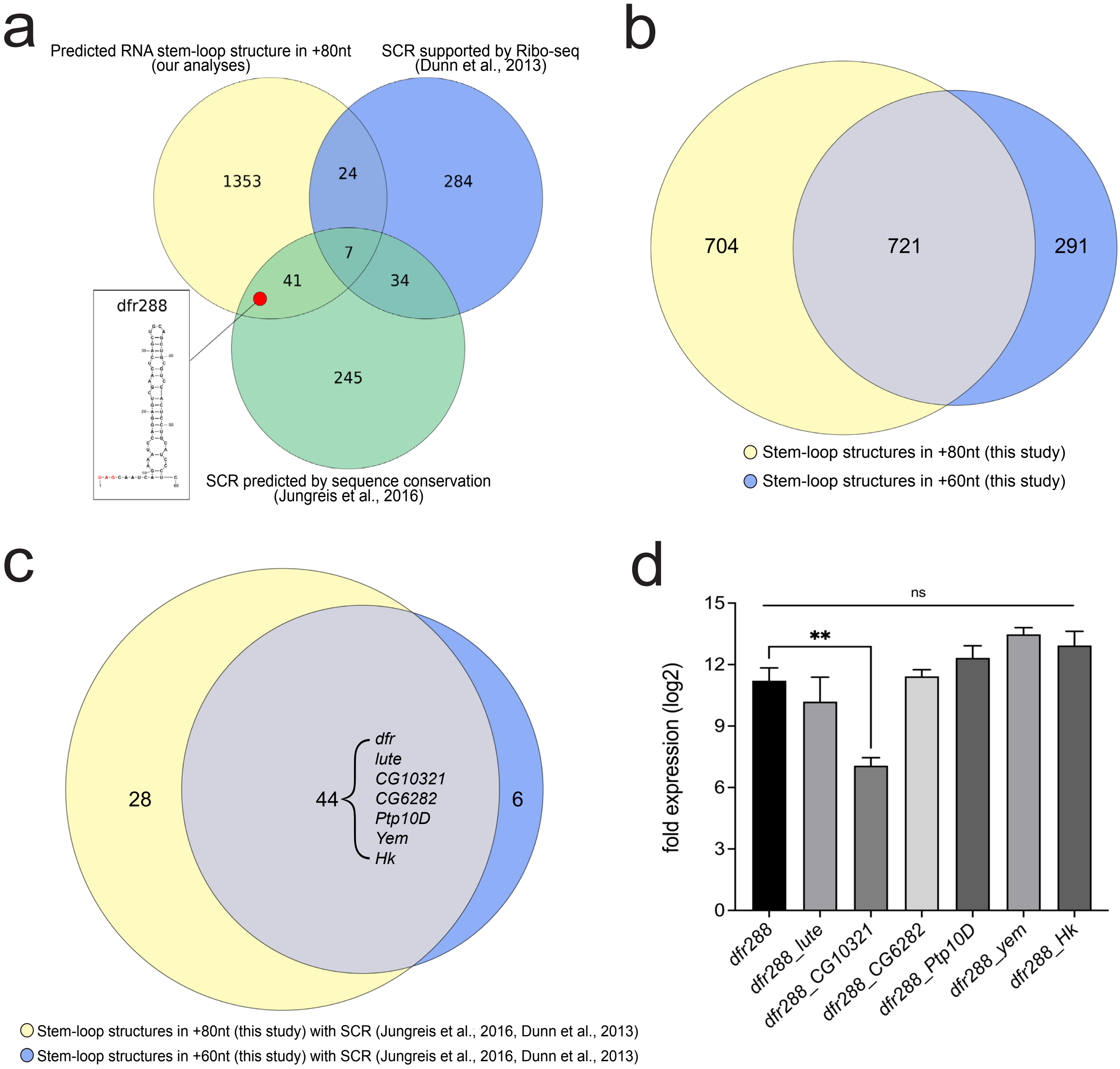


#### Supplementary Fig S5
