## Supplementary Data SD2 for "The mRNA architecture of the termination site primes programmed stop codon readthrough events in *Drosophila*"

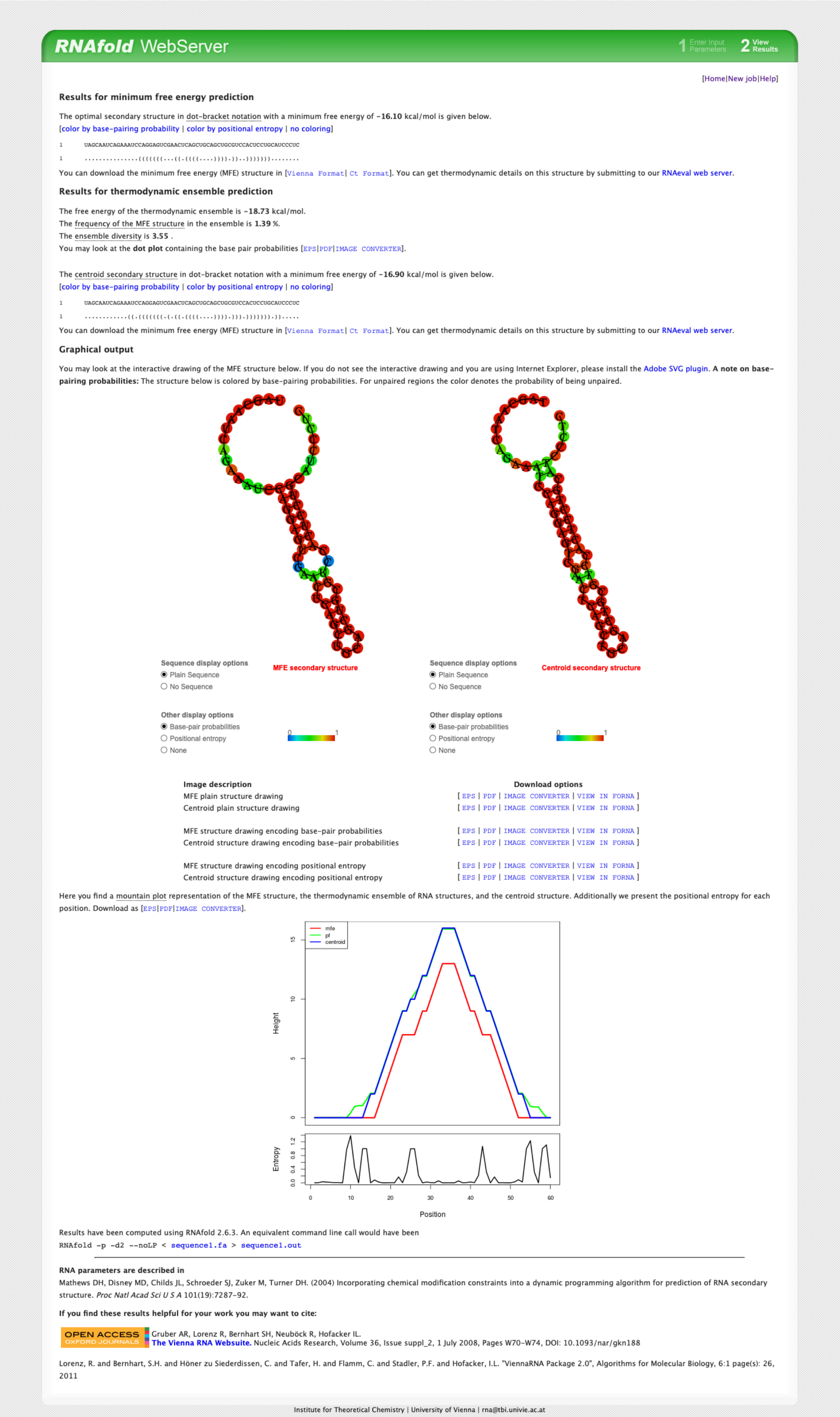

RNAfold WebServer

1 Enter Input Parameters 2 View Results

[Home|New job|Help]

**Results for minimum free energy prediction**

The optimal secondary structure in dot-bracket notation with a minimum free energy of **-16.10** kcal/mol is given below.
[color by base-pairing probability | color by positional entropy | no coloring]

```
1       UAGCAAUCAGAAAUCCAGGAGUCGAACUCAGCUGCAGCUGCGUCCACUCCUGCAUCCCUC
1       ...............(((((((...((.((((....)))).))..)))))))........
```

You can download the minimum free energy (MFE) structure in [Vienna Format| Ct Format]. You can get thermodynamic details on this structure by submitting to our RNAeval web server.

**Results for thermodynamic ensemble prediction**

The free energy of the thermodynamic ensemble is **-18.73** kcal/mol.
The frequency of the MFE structure in the ensemble is **1.39** %.
The ensemble diversity is **3.55** .
You may look at the **dot plot** containing the base pair probabilities [EPS|PDF|IMAGE CONVERTER].

The centroid secondary structure in dot-bracket notation with a minimum free energy of **-16.90** kcal/mol is given below.
[color by base-pairing probability | color by positional entropy | no coloring]

```
1       UAGCAAUCAGAAAUCCAGGAGUCGAACUCAGCUGCAGCUGCGUCCACUCCUGCAUCCCUC
1       ...........((.(((((((.(.((.((((....)))).))).))))))).)).....
```

You can download the minimum free energy (MFE) structure in [Vienna Format| Ct Format]. You can get thermodynamic details on this structure by submitting to our RNAeval web server.

**Graphical output**

You may look at the interactive drawing of the MFE structure below. If you do not see the interactive drawing and you are using Internet Explorer, please install the Adobe SVG plugin. **A note on base-pairing probabilities:** The structure below is colored by base-pairing probabilities. For unpaired regions the color denotes the probability of being unpaired.

MFE secondary structure

Sequence display options
Plain Sequence
No Sequence

Other display options
Base-pair probabilities
Positional entropy
None

0 1

Centroid secondary structure

Sequence display options
Plain Sequence
No Sequence

Other display options
Base-pair probabilities
Positional entropy
None

0 1

| Image description | Download options |
|---|---|
| MFE plain structure drawing | [ EPS \| PDF \| IMAGE CONVERTER \| VIEW IN FORNA ] |
| Centroid plain structure drawing | [ EPS \| PDF \| IMAGE CONVERTER \| VIEW IN FORNA ] |
| MFE structure drawing encoding base-pair probabilities | [ EPS \| PDF \| IMAGE CONVERTER \| VIEW IN FORNA ] |
| Centroid structure drawing encoding base-pair probabilities | [ EPS \| PDF \| IMAGE CONVERTER \| VIEW IN FORNA ] |
| MFE structure drawing encoding positional entropy | [ EPS \| PDF \| IMAGE CONVERTER \| VIEW IN FORNA ] |
| Centroid structure drawing encoding positional entropy | [ EPS \| PDF \| IMAGE CONVERTER \| VIEW IN FORNA ] |

Here you find a mountain plot representation of the MFE structure, the thermodynamic ensemble of RNA structures, and the centroid structure. Additionally we present the positional entropy for each position. Download as [EPS|PDF|IMAGE CONVERTER].



Results have been computed using RNAfold 2.6.3. An equivalent command line call would have been

```
RNAfold -p -d2 --noLP < sequence1.fa > sequence1.out
```

**RNA parameters are described in**

Mathews DH, Disney MD, Childs JL, Schroeder SJ, Zuker M, Turner DH. (2004) Incorporating chemical modification constraints into a dynamic programming algorithm for prediction of RNA secondary structure. *Proc Natl Acad Sci U S A* 101(19):7287-92.

**If you find these results helpful for your work you may want to cite:**

OPEN ACCESS OXFORD JOURNALS Gruber AR, Lorenz R, Bernhart SH, Neuböck R, Hofacker IL.
**The Vienna RNA Websuite.** Nucleic Acids Research, Volume 36, Issue suppl_2, 1 July 2008, Pages W70-W74, DOI: 10.1093/nar/gkn188

Lorenz, R. and Bernhart, S.H. and Höner zu Siederdissen, C. and Tafer, H. and Flamm, C. and Stadler, P.F. and Hofacker, I.L. "ViennaRNA Package 2.0", Algorithms for Molecular Biology, 6:1 page(s): 26, 2011

Institute for Theoretical Chemistry | University of Vienna |

*dfr/vvl* mRNA secondary structure prediction with RNAfold (http://rna.tbi.univie.ac.at//cgi-bin/RNAWebSuite/RNAfold.cgi)

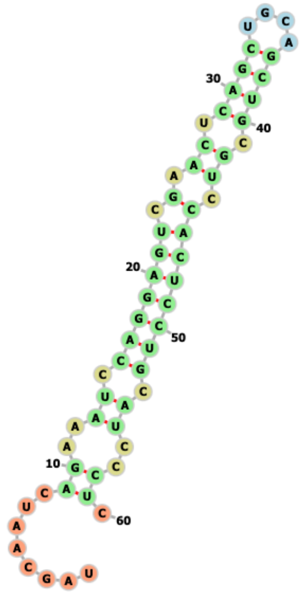


*dfr/vvl* mRNA secondary structure prediction with mxfold2
(https://ws.sato-lab.org/mxfold2/)

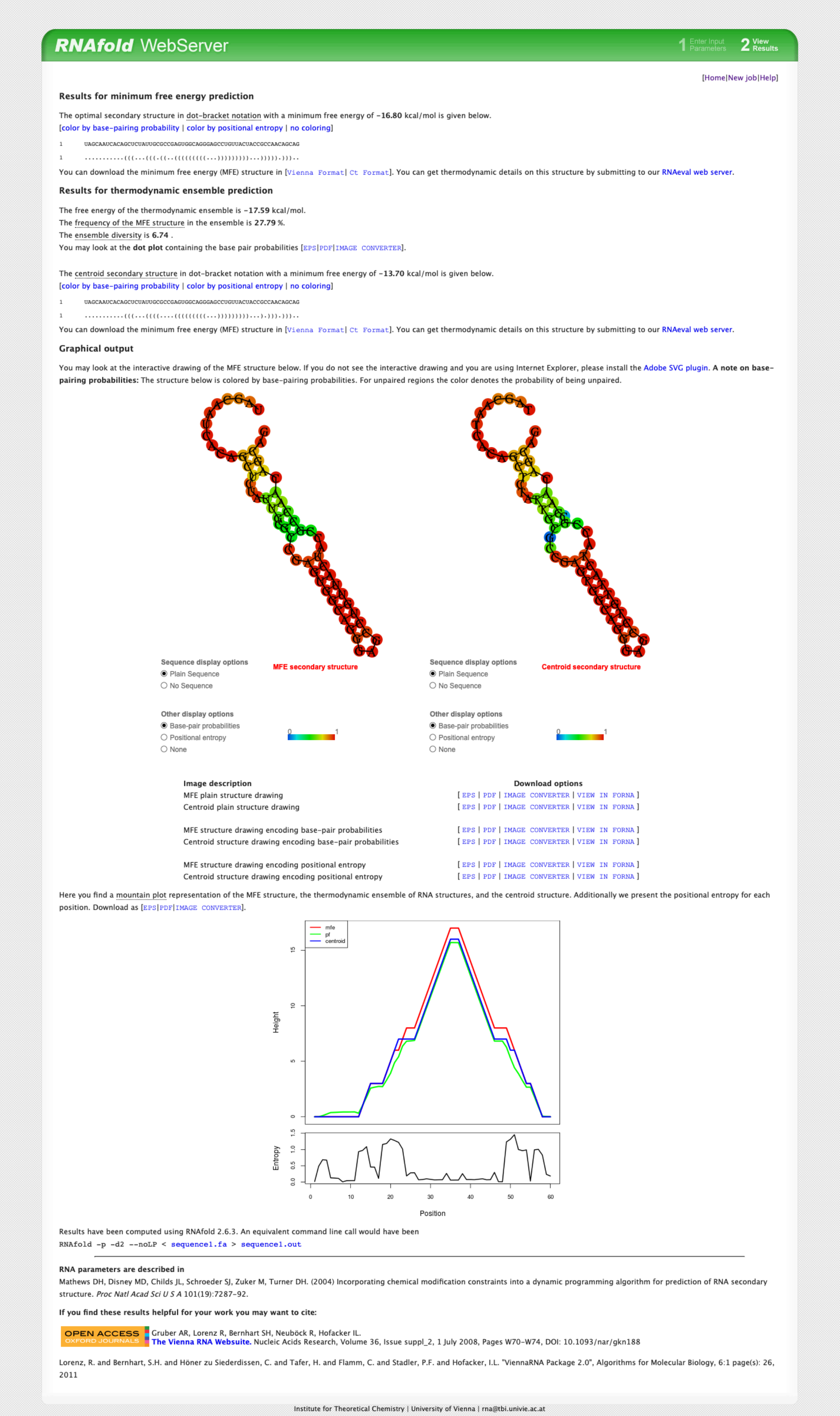

**RNAfold** WebServer — 1 Enter Input Parameters — 2 View Results

[Home|New job|Help]

**Results for minimum free energy prediction**

The optimal secondary structure in dot-bracket notation with a minimum free energy of **-16.80** kcal/mol is given below.
[color by base-pairing probability | color by positional entropy | no coloring]

```
1       UAGCAAUCACAGCUCUAUUGCGCCGAGUGGCAGGGAGCCUGUUACUACCGCCAACAGCAG
1       ...........(((...(((.((..(((((((((...)))))))))...))))).)))..
```

You can download the minimum free energy (MFE) structure in [Vienna Format| Ct Format]. You can get thermodynamic details on this structure by submitting to our RNAeval web server.

**Results for thermodynamic ensemble prediction**

The free energy of the thermodynamic ensemble is **-17.59** kcal/mol.
The frequency of the MFE structure in the ensemble is **27.79** %.
The ensemble diversity is **6.74** .
You may look at the **dot plot** containing the base pair probabilities [EPS|PDF|IMAGE CONVERTER].

The centroid secondary structure in dot-bracket notation with a minimum free energy of **-13.70** kcal/mol is given below.
[color by base-pairing probability | color by positional entropy | no coloring]

```
1       UAGCAAUCACAGCUCUAUUGCGCCGAGUGGCAGGGAGCCUGUUACUACCGCCAACAGCAG
1       ...........(((...((((....(((((((((...)))))))))...).))).)))..
```

You can download the minimum free energy (MFE) structure in [Vienna Format| Ct Format]. You can get thermodynamic details on this structure by submitting to our RNAeval web server.

**Graphical output**

You may look at the interactive drawing of the MFE structure below. If you do not see the interactive drawing and you are using Internet Explorer, please install the Adobe SVG plugin. **A note on base-pairing probabilities:** The structure below is colored by base-pairing probabilities. For unpaired regions the color denotes the probability of being unpaired.

MFE secondary structure — Centroid secondary structure

Sequence display options: Plain Sequence / No Sequence
Other display options: Base-pair probabilities / Positional entropy / None

| Image description | Download options |
|---|---|
| MFE plain structure drawing | [ EPS \| PDF \| IMAGE CONVERTER \| VIEW IN FORNA ] |
| Centroid plain structure drawing | [ EPS \| PDF \| IMAGE CONVERTER \| VIEW IN FORNA ] |
| MFE structure drawing encoding base-pair probabilities | [ EPS \| PDF \| IMAGE CONVERTER \| VIEW IN FORNA ] |
| Centroid structure drawing encoding base-pair probabilities | [ EPS \| PDF \| IMAGE CONVERTER \| VIEW IN FORNA ] |
| MFE structure drawing encoding positional entropy | [ EPS \| PDF \| IMAGE CONVERTER \| VIEW IN FORNA ] |
| Centroid structure drawing encoding positional entropy | [ EPS \| PDF \| IMAGE CONVERTER \| VIEW IN FORNA ] |

Here you find a mountain plot representation of the MFE structure, the thermodynamic ensemble of RNA structures, and the centroid structure. Additionally we present the positional entropy for each position. Download as [EPS|PDF|IMAGE CONVERTER].



Results have been computed using RNAfold 2.6.3. An equivalent command line call would have been
`RNAfold -p -d2 --noLP < sequence1.fa > sequence1.out`

**RNA parameters are described in**
Mathews DH, Disney MD, Childs JL, Schroeder SJ, Zuker M, Turner DH. (2004) Incorporating chemical modification constraints into a dynamic programming algorithm for prediction of RNA secondary structure. *Proc Natl Acad Sci U S A* 101(19):7287-92.

**If you find these results helpful for your work you may want to cite:**

OPEN ACCESS OXFORD JOURNALS — Gruber AR, Lorenz R, Bernhart SH, Neuböck R, Hofacker IL. **The Vienna RNA Websuite.** Nucleic Acids Research, Volume 36, Issue suppl_2, 1 July 2008, Pages W70–W74, DOI: 10.1093/nar/gkn188

Lorenz, R. and Bernhart, S.H. and Höner zu Siederdissen, C. and Tafer, H. and Flamm, C. and Stadler, P.F. and Hofacker, I.L. "ViennaRNA Package 2.0", Algorithms for Molecular Biology, 6:1 page(s): 26, 2011

Institute for Theoretical Chemistry | University of Vienna |

*lute* mRNA secondary structure prediction with RNAfold (http://rna.tbi.univie.ac.at//cgi-bin/RNAWebSuite/RNAfold.cgi)

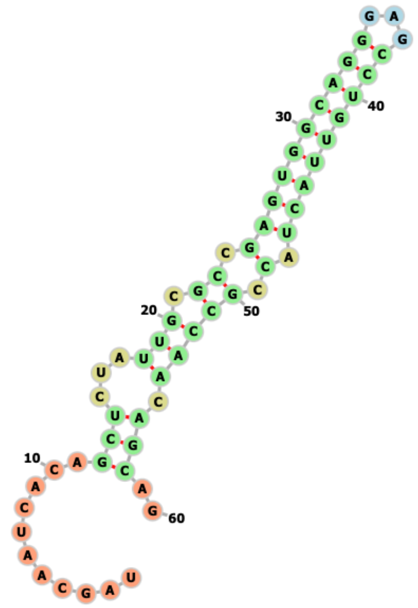


*lute* mRNA secondary structure prediction with mxfold2
(https://ws.sato-lab.org/mxfold2/)

**RNAfold** WebServer

1 Enter Input Parameters  2 View Results

[Home|New job|Help]

### Results for minimum free energy prediction

The optimal secondary structure in dot-bracket notation with a minimum free energy of **-34.10** kcal/mol is given below.
[color by base-pairing probability | color by positional entropy | no coloring]

```
1       UAGCAAUCAGAGUUUGCCGCAGCGGGCGCUCCUGGAAACGGCGAGUGCCUGCUCGGCAUU
1       ..............((((.(((((((((((((((....))).))))))))))))))))..
```

You can download the minimum free energy (MFE) structure in [Vienna Format| Ct Format]. You can get thermodynamic details on this structure by submitting to our RNAeval web server.

### Results for thermodynamic ensemble prediction

The free energy of the thermodynamic ensemble is **-34.35** kcal/mol.
The frequency of the MFE structure in the ensemble is **66.93** %.
The ensemble diversity is **1.07** .
You may look at the **dot plot** containing the base pair probabilities [EPS|PDF|IMAGE CONVERTER].

The centroid secondary structure in dot-bracket notation with a minimum free energy of **-34.10** kcal/mol is given below.
[color by base-pairing probability | color by positional entropy | no coloring]

```
1       UAGCAAUCAGAGUUUGCCGCAGCGGGCGCUCCUGGAAACGGCGAGUGCCUGCUCGGCAUU
1       ..............((((.(((((((((((((((....))).))))))))))))))))..
```

You can download the minimum free energy (MFE) structure in [Vienna Format| Ct Format]. You can get thermodynamic details on this structure by submitting to our RNAeval web server.

### Graphical output

You may look at the interactive drawing of the MFE structure below. If you do not see the interactive drawing and you are using Internet Explorer, please install the Adobe SVG plugin. **A note on base-pairing probabilities:** The structure below is colored by base-pairing probabilities. For unpaired regions the color denotes the probability of being unpaired.

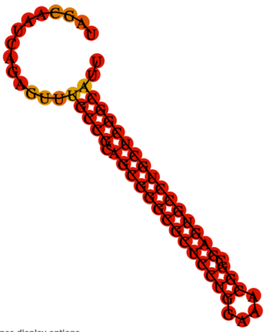

Sequence display options
- Plain Sequence
- No Sequence

**MFE secondary structure**

Other display options
- Base-pair probabilities
- Positional entropy
- None

0 1

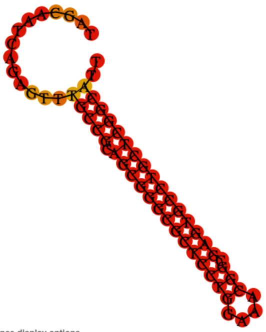

Sequence display options
- Plain Sequence
- No Sequence

**Centroid secondary structure**

Other display options
- Base-pair probabilities
- Positional entropy
- None

0 1

| Image description | Download options |
|---|---|
| MFE plain structure drawing | [ EPS \| PDF \| IMAGE CONVERTER \| VIEW IN FORNA ] |
| Centroid plain structure drawing | [ EPS \| PDF \| IMAGE CONVERTER \| VIEW IN FORNA ] |
| MFE structure drawing encoding base-pair probabilities | [ EPS \| PDF \| IMAGE CONVERTER \| VIEW IN FORNA ] |
| Centroid structure drawing encoding base-pair probabilities | [ EPS \| PDF \| IMAGE CONVERTER \| VIEW IN FORNA ] |
| MFE structure drawing encoding positional entropy | [ EPS \| PDF \| IMAGE CONVERTER \| VIEW IN FORNA ] |
| Centroid structure drawing encoding positional entropy | [ EPS \| PDF \| IMAGE CONVERTER \| VIEW IN FORNA ] |

Here you find a mountain plot representation of the MFE structure, the thermodynamic ensemble of RNA structures, and the centroid structure. Additionally we present the positional entropy for each position. Download as [EPS|PDF|IMAGE CONVERTER].

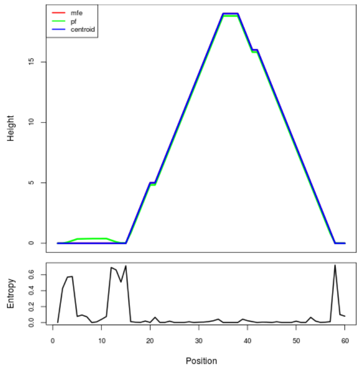


Results have been computed using RNAfold 2.6.3. An equivalent command line call would have been

```
RNAfold -p -d2 --noLP < sequence1.fa > sequence1.out
```

**RNA parameters are described in**

Mathews DH, Disney MD, Childs JL, Schroeder SJ, Zuker M, Turner DH. (2004) Incorporating chemical modification constraints into a dynamic programming algorithm for prediction of RNA secondary structure. *Proc Natl Acad Sci U S A* 101(19):7287-92.

**If you find these results helpful for your work you may want to cite:**

OPEN ACCESS OXFORD JOURNALS Gruber AR, Lorenz R, Bernhart SH, Neuböck R, Hofacker IL.
**The Vienna RNA Websuite.** Nucleic Acids Research, Volume 36, Issue suppl_2, 1 July 2008, Pages W70–W74, DOI: 10.1093/nar/gkn188

Lorenz, R. and Bernhart, S.H. and Höner zu Siederdissen, C. and Tafer, H. and Flamm, C. and Stadler, P.F. and Hofacker, I.L. "ViennaRNA Package 2.0", Algorithms for Molecular Biology, 6:1 page(s): 26, 2011

Institute for Theoretical Chemistry | University of Vienna |

*CG10321* mRNA secondary structure prediction with RNAfold (http://rna.tbi.univie.ac.at//cgi-bin/RNAWebSuite/RNAfold.cgi)

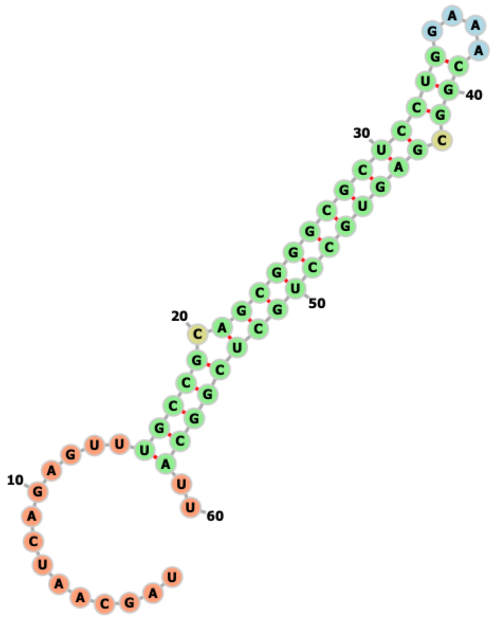


*CG10321* mRNA secondary structure prediction with mxfold2
(https://ws.sato-lab.org/mxfold2/)

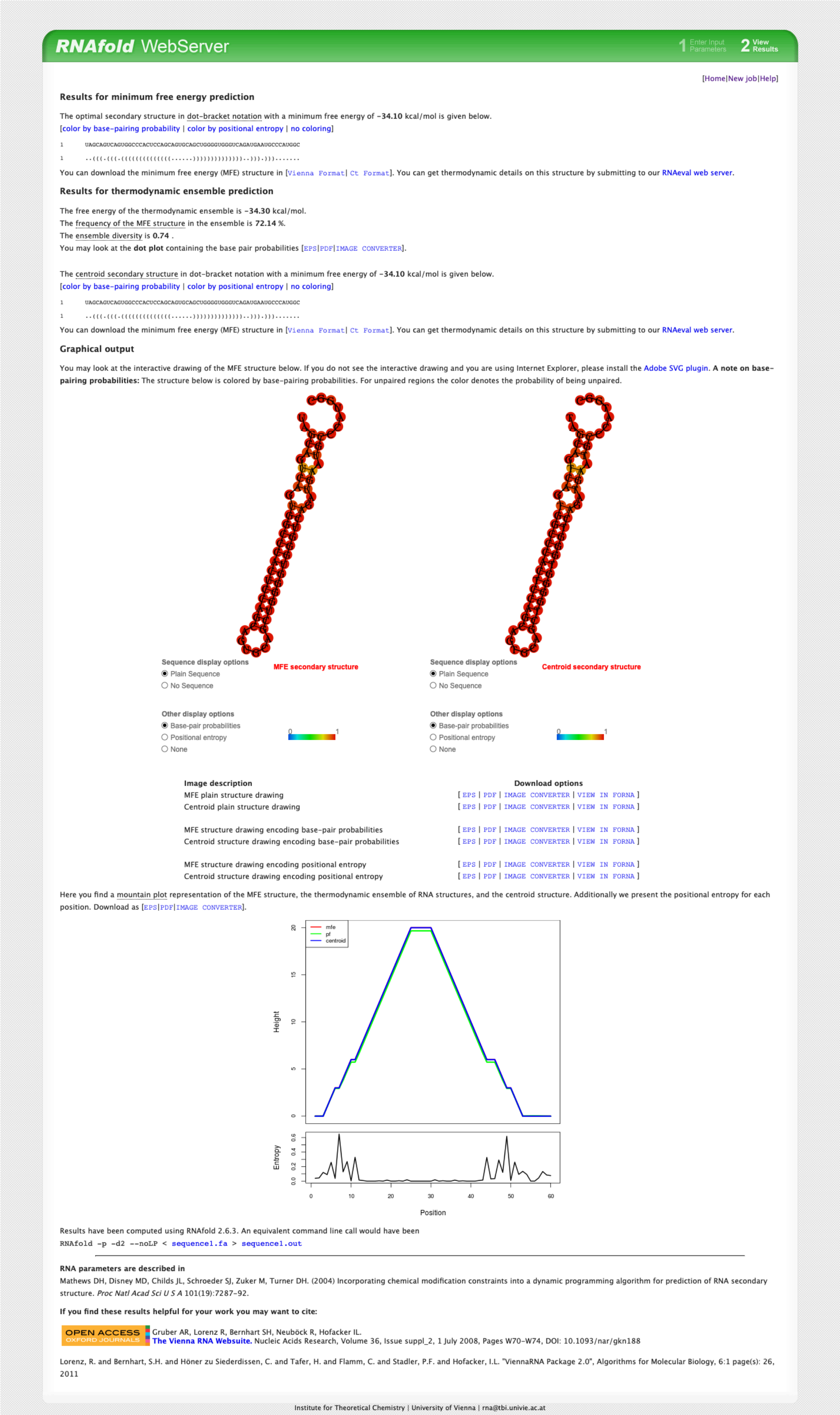

RNAfold WebServer

1 Enter Input Parameters 2 View Results

[Home|New job|Help]

**Results for minimum free energy prediction**

The optimal secondary structure in dot-bracket notation with a minimum free energy of **-34.10** kcal/mol is given below.
[color by base-pairing probability | color by positional entropy | no coloring]

```
1       UAGCAGUCAGUGGCCCACUCCAGCAGUGCAGCUGGGGUGGGUCAGAUGAAUGCCCAUGGC
1       ..(((.(((.((((((((((((((......))))))))))))))..))).))).......
```

You can download the minimum free energy (MFE) structure in [Vienna Format| Ct Format]. You can get thermodynamic details on this structure by submitting to our RNAeval web server.

**Results for thermodynamic ensemble prediction**

The free energy of the thermodynamic ensemble is **-34.30** kcal/mol.
The frequency of the MFE structure in the ensemble is **72.14** %.
The ensemble diversity is **0.74** .
You may look at the **dot plot** containing the base pair probabilities [EPS|PDF|IMAGE CONVERTER].

The centroid secondary structure in dot-bracket notation with a minimum free energy of **-34.10** kcal/mol is given below.
[color by base-pairing probability | color by positional entropy | no coloring]

```
1       UAGCAGUCAGUGGCCCACUCCAGCAGUGCAGCUGGGGUGGGUCAGAUGAAUGCCCAUGGC
1       ..(((.(((.((((((((((((((......))))))))))))))..))).))).......
```

You can download the minimum free energy (MFE) structure in [Vienna Format| Ct Format]. You can get thermodynamic details on this structure by submitting to our RNAeval web server.

**Graphical output**

You may look at the interactive drawing of the MFE structure below. If you do not see the interactive drawing and you are using Internet Explorer, please install the Adobe SVG plugin. **A note on base-pairing probabilities:** The structure below is colored by base-pairing probabilities. For unpaired regions the color denotes the probability of being unpaired.



| Image description | Download options |
| --- | --- |
| MFE plain structure drawing | [ EPS \| PDF \| IMAGE CONVERTER \| VIEW IN FORNA ] |
| Centroid plain structure drawing | [ EPS \| PDF \| IMAGE CONVERTER \| VIEW IN FORNA ] |
| MFE structure drawing encoding base-pair probabilities | [ EPS \| PDF \| IMAGE CONVERTER \| VIEW IN FORNA ] |
| Centroid structure drawing encoding base-pair probabilities | [ EPS \| PDF \| IMAGE CONVERTER \| VIEW IN FORNA ] |
| MFE structure drawing encoding positional entropy | [ EPS \| PDF \| IMAGE CONVERTER \| VIEW IN FORNA ] |
| Centroid structure drawing encoding positional entropy | [ EPS \| PDF \| IMAGE CONVERTER \| VIEW IN FORNA ] |

Here you find a mountain plot representation of the MFE structure, the thermodynamic ensemble of RNA structures, and the centroid structure. Additionally we present the positional entropy for each position. Download as [EPS|PDF|IMAGE CONVERTER].



Results have been computed using RNAfold 2.6.3. An equivalent command line call would have been

```
RNAfold -p -d2 --noLP < sequence1.fa > sequence1.out
```

**RNA parameters are described in**


Mathews DH, Disney MD, Childs JL, Schroeder SJ, Zuker M, Turner DH. (2004) Incorporating chemical modification constraints into a dynamic programming algorithm for prediction of RNA secondary structure. *Proc Natl Acad Sci U S A* 101(19):7287-92.


**If you find these results helpful for your work you may want to cite:**


OPEN ACCESS OXFORD JOURNALS Gruber AR, Lorenz R, Bernhart SH, Neuböck R, Hofacker IL. **The Vienna RNA Websuite.** Nucleic Acids Research, Volume 36, Issue suppl_2, 1 July 2008, Pages W70-W74, DOI: 10.1093/nar/gkn188

Lorenz, R. and Bernhart, S.H. and Höner zu Siederdissen, C. and Tafer, H. and Flamm, C. and Stadler, P.F. and Hofacker, I.L. "ViennaRNA Package 2.0", Algorithms for Molecular Biology, 6:1 page(s): 26, 2011


Institute for Theoretical Chemistry | University of Vienna |

*CG6282* mRNA secondary structure prediction with RNAfold (http://rna.tbi.univie.ac.at//cgi-bin/RNAWebSuite/RNAfold.cgi)

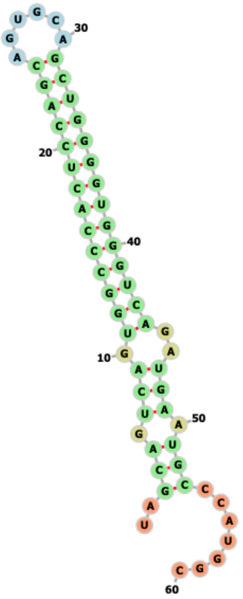


*CG6282* mRNA secondary structure prediction with mxfold2 (https://ws.sato-lab.org/mxfold2/)

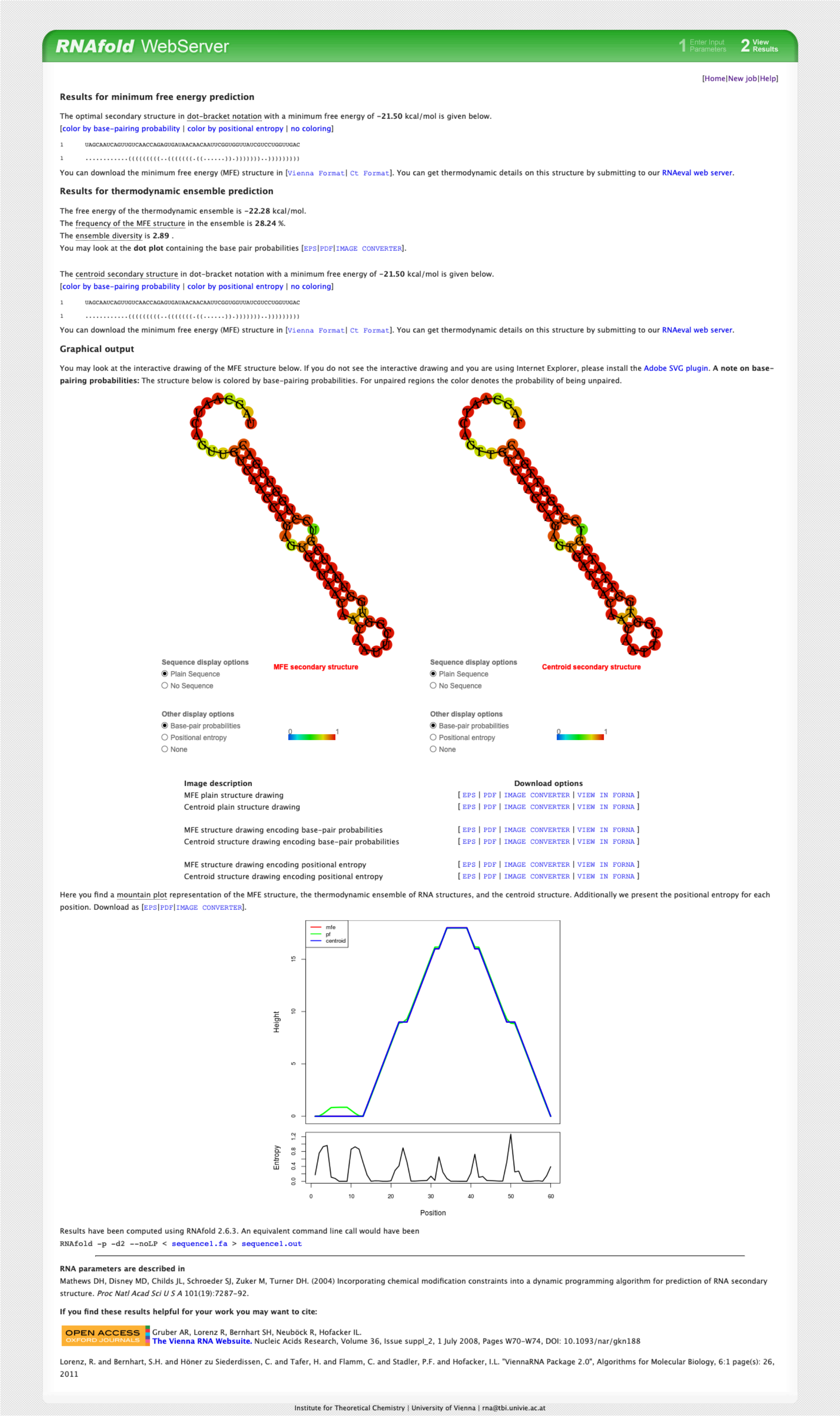

**RNAfold** WebServer

1 Enter Input Parameters 2 View Results

[Home|New job|Help]

**Results for minimum free energy prediction**

The optimal secondary structure in dot-bracket notation with a minimum free energy of **-21.50** kcal/mol is given below.
[color by base-pairing probability | color by positional entropy | no coloring]

```
1       UAGCAAUCAGUUGUCAACCAGAGUGAUAACAACAAUUCGGUGGUUAUCGUCCUGGUUGAC
1       ............(((((((((..(((((((.((......)).)))))))..)))))))))
```

You can download the minimum free energy (MFE) structure in [Vienna Format| Ct Format]. You can get thermodynamic details on this structure by submitting to our RNAeval web server.

**Results for thermodynamic ensemble prediction**

The free energy of the thermodynamic ensemble is **-22.28** kcal/mol.
The frequency of the MFE structure in the ensemble is **28.24** %.
The ensemble diversity is **2.89** .
You may look at the **dot plot** containing the base pair probabilities [EPS|PDF|IMAGE CONVERTER].

The centroid secondary structure in dot-bracket notation with a minimum free energy of **-21.50** kcal/mol is given below.
[color by base-pairing probability | color by positional entropy | no coloring]

```
1       UAGCAAUCAGUUGUCAACCAGAGUGAUAACAACAAUUCGGUGGUUAUCGUCCUGGUUGAC
1       ............(((((((((..(((((((.((......)).)))))))..)))))))))
```

You can download the minimum free energy (MFE) structure in [Vienna Format| Ct Format]. You can get thermodynamic details on this structure by submitting to our RNAeval web server.

**Graphical output**

You may look at the interactive drawing of the MFE structure below. If you do not see the interactive drawing and you are using Internet Explorer, please install the Adobe SVG plugin. **A note on base-pairing probabilities:** The structure below is colored by base-pairing probabilities. For unpaired regions the color denotes the probability of being unpaired.



| Image description | Download options |
|---|---|
| MFE plain structure drawing | [ EPS \| PDF \| IMAGE CONVERTER \| VIEW IN FORNA ] |
| Centroid plain structure drawing | [ EPS \| PDF \| IMAGE CONVERTER \| VIEW IN FORNA ] |
| MFE structure drawing encoding base-pair probabilities | [ EPS \| PDF \| IMAGE CONVERTER \| VIEW IN FORNA ] |
| Centroid structure drawing encoding base-pair probabilities | [ EPS \| PDF \| IMAGE CONVERTER \| VIEW IN FORNA ] |
| MFE structure drawing encoding positional entropy | [ EPS \| PDF \| IMAGE CONVERTER \| VIEW IN FORNA ] |
| Centroid structure drawing encoding positional entropy | [ EPS \| PDF \| IMAGE CONVERTER \| VIEW IN FORNA ] |

Here you find a mountain plot representation of the MFE structure, the thermodynamic ensemble of RNA structures, and the centroid structure. Additionally we present the positional entropy for each position. Download as [EPS|PDF|IMAGE CONVERTER].



Results have been computed using RNAfold 2.6.3. An equivalent command line call would have been

```
RNAfold -p -d2 --noLP < sequence1.fa > sequence1.out
```

**RNA parameters are described in**

Mathews DH, Disney MD, Childs JL, Schroeder SJ, Zuker M, Turner DH. (2004) Incorporating chemical modification constraints into a dynamic programming algorithm for prediction of RNA secondary structure. *Proc Natl Acad Sci U S A* 101(19):7287-92.

**If you find these results helpful for your work you may want to cite:**

OPEN ACCESS OXFORD JOURNALS Gruber AR, Lorenz R, Bernhart SH, Neuböck R, Hofacker IL.
**The Vienna RNA Websuite.** Nucleic Acids Research, Volume 36, Issue suppl_2, 1 July 2008, Pages W70-W74, DOI: 10.1093/nar/gkn188

Lorenz, R. and Bernhart, S.H. and Höner zu Siederdissen, C. and Tafer, H. and Flamm, C. and Stadler, P.F. and Hofacker, I.L. "ViennaRNA Package 2.0", Algorithms for Molecular Biology, 6:1 page(s): 26, 2011

Institute for Theoretical Chemistry | University of Vienna |

*Ptp10D* mRNA secondary structure prediction with RNAfold (http://rna.tbi.univie.ac.at//cgi-bin/RNAWebSuite/RNAfold.cgi)

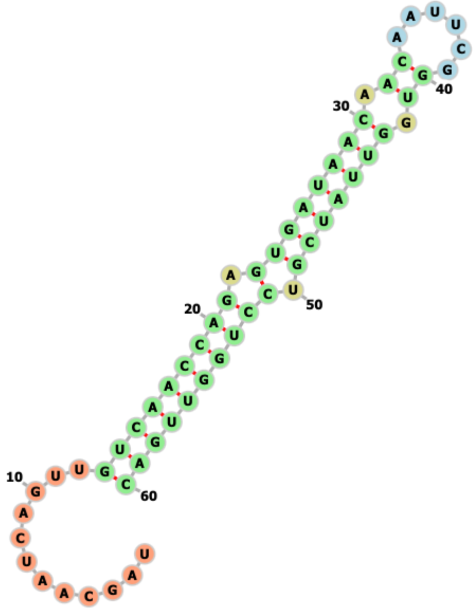


*Ptp10D* mRNA secondary structure prediction with mxfold2 (https://ws.sato-lab.org/mxfold2/)

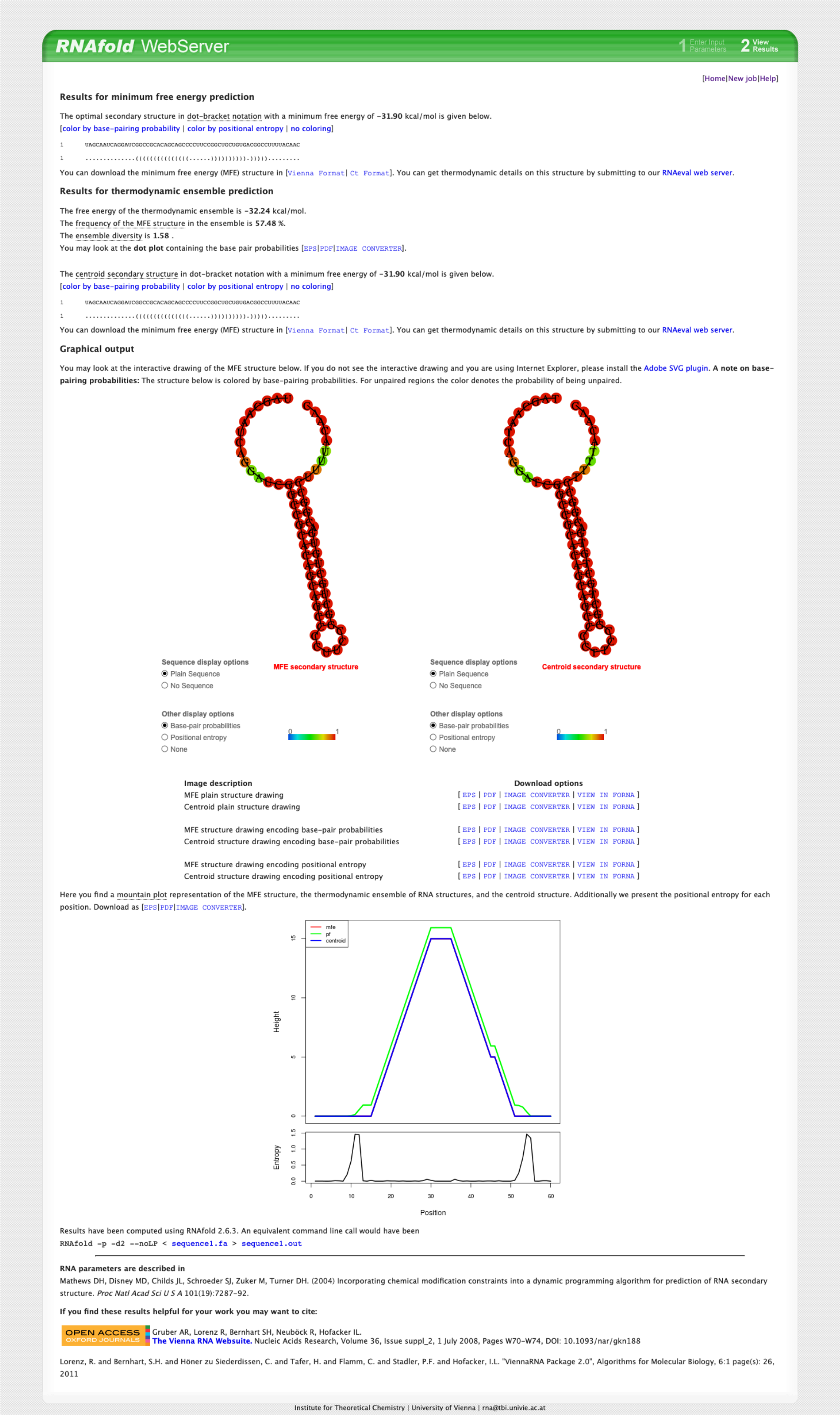

RNAfold WebServer

1 Enter Input Parameters 2 View Results

[Home|New job|Help]

**Results for minimum free energy prediction**

The optimal secondary structure in dot-bracket notation with a minimum free energy of **-31.90** kcal/mol is given below.
[color by base-pairing probability | color by positional entropy | no coloring]

```
1      UAGCAAUCAGGAUCGGCCGCACAGCAGCCCCUUCCGGCUGCUGUGACGGCCUUUUACAAC
1      ..............((((((((((((((......))))))))))).)))))........ 
```

You can download the minimum free energy (MFE) structure in [Vienna Format| Ct Format]. You can get thermodynamic details on this structure by submitting to our RNAeval web server.

**Results for thermodynamic ensemble prediction**

The free energy of the thermodynamic ensemble is **-32.24** kcal/mol.
The frequency of the MFE structure in the ensemble is **57.48** %.
The ensemble diversity is **1.58** .
You may look at the **dot plot** containing the base pair probabilities [EPS|PDF|IMAGE CONVERTER].

The centroid secondary structure in dot-bracket notation with a minimum free energy of **-31.90** kcal/mol is given below.
[color by base-pairing probability | color by positional entropy | no coloring]

```
1      UAGCAAUCAGGAUCGGCCGCACAGCAGCCCCUUCCGGCUGCUGUGACGGCCUUUUACAAC
1      ..............((((((((((((((......))))))))))).)))))........ 
```

You can download the minimum free energy (MFE) structure in [Vienna Format| Ct Format]. You can get thermodynamic details on this structure by submitting to our RNAeval web server.

**Graphical output**

You may look at the interactive drawing of the MFE structure below. If you do not see the interactive drawing and you are using Internet Explorer, please install the Adobe SVG plugin. **A note on base-pairing probabilities:** The structure below is colored by base-pairing probabilities. For unpaired regions the color denotes the probability of being unpaired.

Sequence display options
Plain Sequence
No Sequence

MFE secondary structure

Other display options
Base-pair probabilities
Positional entropy
None

0 1

Sequence display options
Plain Sequence
No Sequence

Centroid secondary structure

Other display options
Base-pair probabilities
Positional entropy
None

0 1

| Image description | Download options |
|---|---|
| MFE plain structure drawing | [ EPS \| PDF \| IMAGE CONVERTER \| VIEW IN FORNA ] |
| Centroid plain structure drawing | [ EPS \| PDF \| IMAGE CONVERTER \| VIEW IN FORNA ] |
| MFE structure drawing encoding base-pair probabilities | [ EPS \| PDF \| IMAGE CONVERTER \| VIEW IN FORNA ] |
| Centroid structure drawing encoding base-pair probabilities | [ EPS \| PDF \| IMAGE CONVERTER \| VIEW IN FORNA ] |
| MFE structure drawing encoding positional entropy | [ EPS \| PDF \| IMAGE CONVERTER \| VIEW IN FORNA ] |
| Centroid structure drawing encoding positional entropy | [ EPS \| PDF \| IMAGE CONVERTER \| VIEW IN FORNA ] |

Here you find a mountain plot representation of the MFE structure, the thermodynamic ensemble of RNA structures, and the centroid structure. Additionally we present the positional entropy for each position. Download as [EPS|PDF|IMAGE CONVERTER].



Results have been computed using RNAfold 2.6.3. An equivalent command line call would have been

```
RNAfold -p -d2 --noLP < sequence1.fa > sequence1.out
```

**RNA parameters are described in**
Mathews DH, Disney MD, Childs JL, Schroeder SJ, Zuker M, Turner DH. (2004) Incorporating chemical modification constraints into a dynamic programming algorithm for prediction of RNA secondary structure. *Proc Natl Acad Sci U S A* 101(19):7287-92.

**If you find these results helpful for your work you may want to cite:**

OPEN ACCESS OXFORD JOURNALS Gruber AR, Lorenz R, Bernhart SH, Neuböck R, Hofacker IL.
**The Vienna RNA Websuite.** Nucleic Acids Research, Volume 36, Issue suppl_2, 1 July 2008, Pages W70-W74, DOI: 10.1093/nar/gkn188

Lorenz, R. and Bernhart, S.H. and Höner zu Siederdissen, C. and Tafer, H. and Flamm, C. and Stadler, P.F. and Hofacker, I.L. "ViennaRNA Package 2.0", Algorithms for Molecular Biology, 6:1 page(s): 26, 2011

Institute for Theoretical Chemistry | University of Vienna |

*Yem* mRNA secondary structure prediction with RNAfold (http://rna.tbi.univie.ac.at//cgi-bin/RNAWebSuite/RNAfold.cgi)

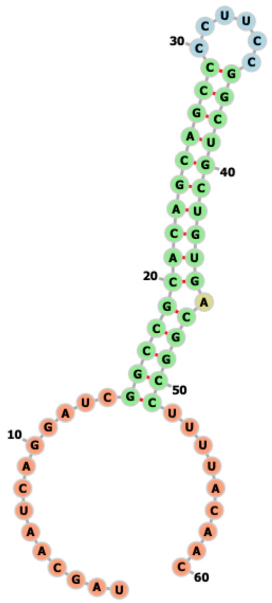


*Yem* mRNA secondary structure prediction with mxfold2 (https://ws.sato-lab.org/mxfold2/)

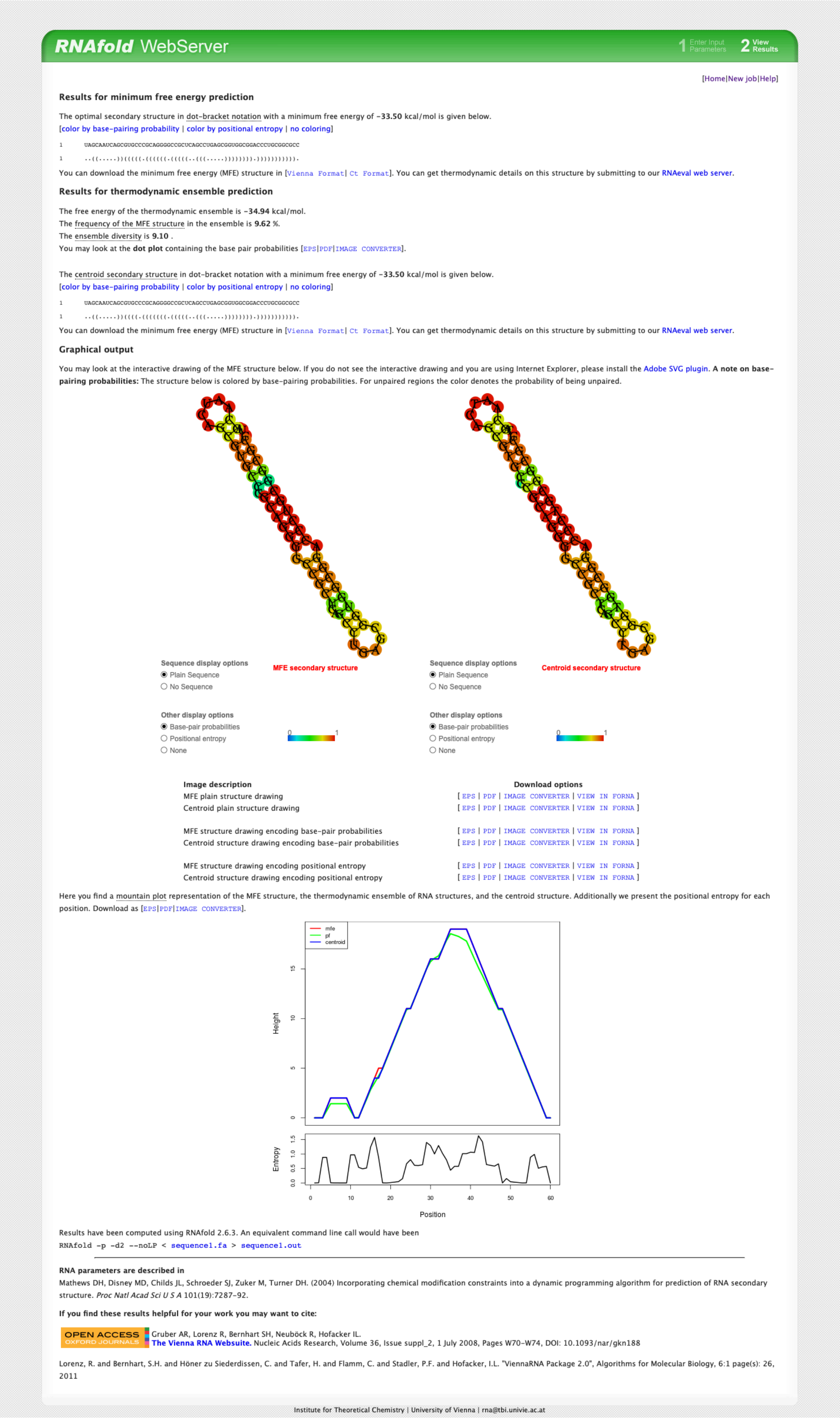

RNAfold WebServer

1 Enter Input Parameters 2 View Results

[Home|New job|Help]

**Results for minimum free energy prediction**

The optimal secondary structure in dot-bracket notation with a minimum free energy of **-33.50** kcal/mol is given below.
[color by base-pairing probability | color by positional entropy | no coloring]

```
1       UAGCAAUCAGCGUGCCCGCAGGGGCCGCUCAGCCUGAGCGGUGGCGGACCCUGCGGCGCC
1       ..((.....))(((((.((((((.(((((..(((.....)))))))).))))))))))).
```

You can download the minimum free energy (MFE) structure in [Vienna Format| Ct Format]. You can get thermodynamic details on this structure by submitting to our RNAeval web server.

**Results for thermodynamic ensemble prediction**

The free energy of the thermodynamic ensemble is **-34.94** kcal/mol.
The frequency of the MFE structure in the ensemble is **9.62** %.
The ensemble diversity is **9.10** .
You may look at the **dot plot** containing the base pair probabilities [EPS|PDF|IMAGE CONVERTER].

The centroid secondary structure in dot-bracket notation with a minimum free energy of **-33.50** kcal/mol is given below.
[color by base-pairing probability | color by positional entropy | no coloring]

```
1       UAGCAAUCAGCGUGCCCGCAGGGGCCGCUCAGCCUGAGCGGUGGCGGACCCUGCGGCGCC
1       ..((.....))((((.(((((((.(((((..(((.....)))))))).))))))))))).
```

You can download the minimum free energy (MFE) structure in [Vienna Format| Ct Format]. You can get thermodynamic details on this structure by submitting to our RNAeval web server.

**Graphical output**

You may look at the interactive drawing of the MFE structure below. If you do not see the interactive drawing and you are using Internet Explorer, please install the Adobe SVG plugin. **A note on base-pairing probabilities:** The structure below is colored by base-pairing probabilities. For unpaired regions the color denotes the probability of being unpaired.



Sequence display options
- Plain Sequence
- No Sequence

Other display options
- Base-pair probabilities
- Positional entropy
- None

| Image description | Download options |
|---|---|
| MFE plain structure drawing | [ EPS \| PDF \| IMAGE CONVERTER \| VIEW IN FORNA ] |
| Centroid plain structure drawing | [ EPS \| PDF \| IMAGE CONVERTER \| VIEW IN FORNA ] |
| MFE structure drawing encoding base-pair probabilities | [ EPS \| PDF \| IMAGE CONVERTER \| VIEW IN FORNA ] |
| Centroid structure drawing encoding base-pair probabilities | [ EPS \| PDF \| IMAGE CONVERTER \| VIEW IN FORNA ] |
| MFE structure drawing encoding positional entropy | [ EPS \| PDF \| IMAGE CONVERTER \| VIEW IN FORNA ] |
| Centroid structure drawing encoding positional entropy | [ EPS \| PDF \| IMAGE CONVERTER \| VIEW IN FORNA ] |

Here you find a mountain plot representation of the MFE structure, the thermodynamic ensemble of RNA structures, and the centroid structure. Additionally we present the positional entropy for each position. Download as [EPS|PDF|IMAGE CONVERTER].



Results have been computed using RNAfold 2.6.3. An equivalent command line call would have been
`RNAfold -p -d2 --noLP < sequence1.fa > sequence1.out`

**RNA parameters are described in**
Mathews DH, Disney MD, Childs JL, Schroeder SJ, Zuker M, Turner DH. (2004) Incorporating chemical modification constraints into a dynamic programming algorithm for prediction of RNA secondary structure. *Proc Natl Acad Sci U S A* 101(19):7287-92.

**If you find these results helpful for your work you may want to cite:**

OPEN ACCESS OXFORD JOURNALS
Gruber AR, Lorenz R, Bernhart SH, Neuböck R, Hofacker IL.
**The Vienna RNA Websuite.** Nucleic Acids Research, Volume 36, Issue suppl_2, 1 July 2008, Pages W70-W74, DOI: 10.1093/nar/gkn188

Lorenz, R. and Bernhart, S.H. and Höner zu Siederdissen, C. and Tafer, H. and Flamm, C. and Stadler, P.F. and Hofacker, I.L. "ViennaRNA Package 2.0", Algorithms for Molecular Biology, 6:1 page(s): 26, 2011

Institute for Theoretical Chemistry | University of Vienna |

*Hk* mRNA secondary structure prediction with RNAfold (http://rna.tbi.univie.ac.at//cgi-bin/RNAWebSuite/RNAfold.cgi)

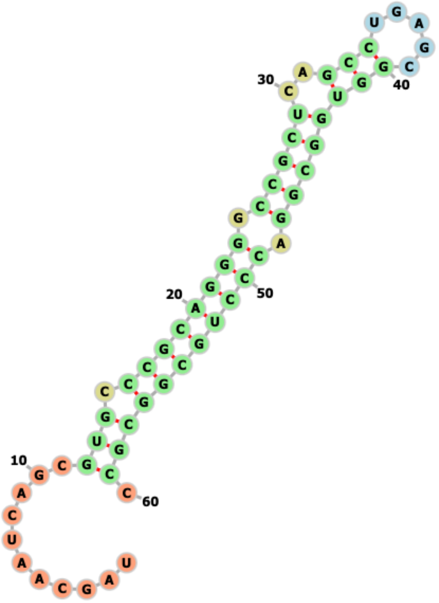


*Hk* mRNA secondary structure prediction with mxfold2 (https://ws.sato-lab.org/mxfold2/)
